# A host AAA-ATPase exhibits bacteriolytic activity for clearance of microbial infection

**DOI:** 10.1101/2023.07.18.549519

**Authors:** Sourav Ghosh, Suvapriya Roy, Navin Baid, Udit Kumar Das, Sumit Rakshit, Paulomi Sanghavi, Dipasree Hajra, Sneha Menon, Mohammad Sahil, Sudipti Shaw, Raju S Rajmani, Harikrishna Adicherla, Jagannath Mondal, Dipshikha Chakravortty, Roop Mallik, Anirban Banerjee

## Abstract

An array of host cytosol guarding factors impede bacterial proliferation and preserve cellular sterility. Amongst them, proteasomal degradation of ubiquitinated pathogens has emerged as a critical mechanism for ensuring cytosolic sanctity. We wondered how proteasomes, with their small size and inability to extract membrane-bound proteins, can eradicate pathogens. Here, we unveil a unique strategy, wherein VCP/p97, a host AAA-ATPase, eliminates pathogens by exerting mechanical force that physically unfolds and pulls out ubiquitinated proteins from bacterial membrane. Combining a single-molecule approach along with molecular dynamic simulation and *in-vitro* reconstitution, we demonstrate that protein extraction by p97 causes extensive membrane lysis and release of cytosolic contents from phylogenetically diverse microbes. Additionally, in an *in-vivo* mouse sepsis model, this segregase-dependent bactericidal effect of p97 abrogated microbial proliferation in host tissues. Overall, we discovered a distinct innate antimicrobial function of p97, that protects the host against lethal bacterial infections.

**One Sentence Summary:** A host AAA-ATPase exhibits bacteriolytic activity.

## Introduction

Cell-autonomous immunity consists of a set of evolutionarily conserved, robust self-defense strategies that metazoans employ to guard their cytosol from pathogenic invasion. Pathogens deploy pore-forming toxins or needle-like secretion systems to catastrophically damage the endosomal membrane, triggering cytosol invasion^1, 2, 3^. To counteract such fatality, the inheritance of various guarding factors is necessary in both immune and non-immune cells. Eukaryotic host has evolved intricate layers of cytosolic pathogen sensing strategies encompassing various Pattern Recognition Receptors (PRRs) which identify diverse Pathogen-associated molecular patterns (PAMPs) such as nucleic acids, lipids, carbohydrates, and peptides^4, 5^. Upon sensing, the host triggers cytosolic effector molecules that participate in pathogen clearance. Members of GBP family (Guanylate binding proteins) majorly exhibit antimicrobial property by targeting the lipopolysaccharides on the Gram-negative bacterial surface^6, 7^. They also form a signalling platform for the initiation of caspase-4-dependent pyroptosis^8, 9, 10^. Additionally, GBPs also permeabilize pathogen’s outer membrane and combine with APOL3 (Apolipoprotein L3), which has detergent-like activity to lyse the bacterial inner membrane^7, 11, 12, 13, 14^. This fine synchronization between a plethora of sensors and executioners forms the decisive fail-safe barrier against a variety of cytosol-invading pathogens. However, these mechanisms are predominantly directed against Gram-negative pathogens. A large number of microbes irrespective of their Gram-origin on the other hand get ubiquitinated for subsequent elimination. Ubiquitination, a cellular homeostatic process^15^, also acts as a significant innate immune barrier in preserving cytosolic sterility. Some E3 Ub-ligases ubiquitinate proteinaceous or non-proteinaceous moieties on microbial surface to drive them either towards autophagy or proteasomal disintegration^9, 12, 16, 17, 18, 19, 20^. While degradation of ubiquitinated pathogens by the autophagy pathway is well elucidated, mechanistic details of proteasomal degradation remained contentious. In general, substrates processed by the proteasome are forced through its barrel, where they are first dismantled and subsequently degraded. However, it is quite unclear how a proteasome can devour an entire bacterium due to the significant difference in size between them (proteasomal entry portal diameter ∼10-20 nm^21^; bacterial size ∼1.5-2 µm). Moreover, bacterial surface proteins that are ubiquitinated, are either cell wall or membrane-bound, posing a challenge for the proteasome to extract them for proteolysis. We therefore speculated that auxiliary host extractors are involved in the perceived antibacterial activity of proteasomes.

AAA-ATPase family proteins are known to be notable segregators in all life forms. Remarkably, these molecular machines are structurally highly conserved and participate in diverse cellular activities ranging from cellular homeostasis to organellar biogenesis^22^. A combination of walker A and B motifs present in these proteins convert chemical energy from ATP hydrolysis into mechanical force to execute its functions^23, 24^. In the host, most of the AAA-ATPases lack proteolytic activity and a major subset of them, direct unfolded substrates towards the proteasomal machinery^25^. In this study, we screened various AAA-ATPases to discover the bacteriolytic role of VCP/p97. Quite remarkably, we show that the forceful extraction of ubiquitinated proteins by VCP/p97 disintegrates the membranes of phylogenetically diverse pathogens leading to their eradication.

### Discovery of VCP/p97 as an antibacterial AAA-ATPase

We first examined whether various host AAA-ATPases, such as p97, PSMC6 (6^th^ subunit of the 19S ATPase complex), MDN1 (Midasin), VPS4 (Vacuolar protein sorting-associated protein), NVL (Nuclear VCP like) and ATAD1 (ATPase family AAA domain containing 1) associate with intracellular SPN (*Streptococcus pneumoniae*). Amongst these, p97 and PSMC6 significantly localized to cell invaded SPN (∼28 and 23%, respectively), while MDN1 (∼8%), VPS4 (∼12%), NVL (∼5%) and ATAD1 (∼4%) showed minimal association (Fig.1A, S1A-F). In addition, p97 co-precipitated with PspA (Pneumococcal surface protein A, reported to be ubiquitinated^12^), whereas other AAA-ATPases displayed little or no interaction (Fig. 1B). Further SPN mutant lacking ubiquitinable surface proteins (PspA and BgaA (β-galactosidase)) showed ∼2.3-fold reduced association with p97 (Fig. S2A). This led us to investigate the consequence of bacterial targeting by p97. Following downregulation of p97’s expression using siRNA, we observed ∼1.8-fold increase in SPN’s intracellular burden (Fig. 1C, S3A). Additionally, burden of two other intracellular pathogens, STm (*Salmonella typhimurium)* and GAS (Group A *Streptococcus*), increased by ∼1.5-2-fold, after reduction of p97’s expression, but not with PSMC6, hinting at a generic guarding tactic employed by p97 against phylogenetically diverse pathogens (Fig. 1C, S3B). Moreover, targeted inhibition of p97’s activity using DBeQ and NMS-873 (at concentrations non-inhibitory to bacterial growth and non-lethal to host cells (Fig. S4A-H) led to elevated bacterial burden inside host cells (Fig. 1D).

**Figure 1.**
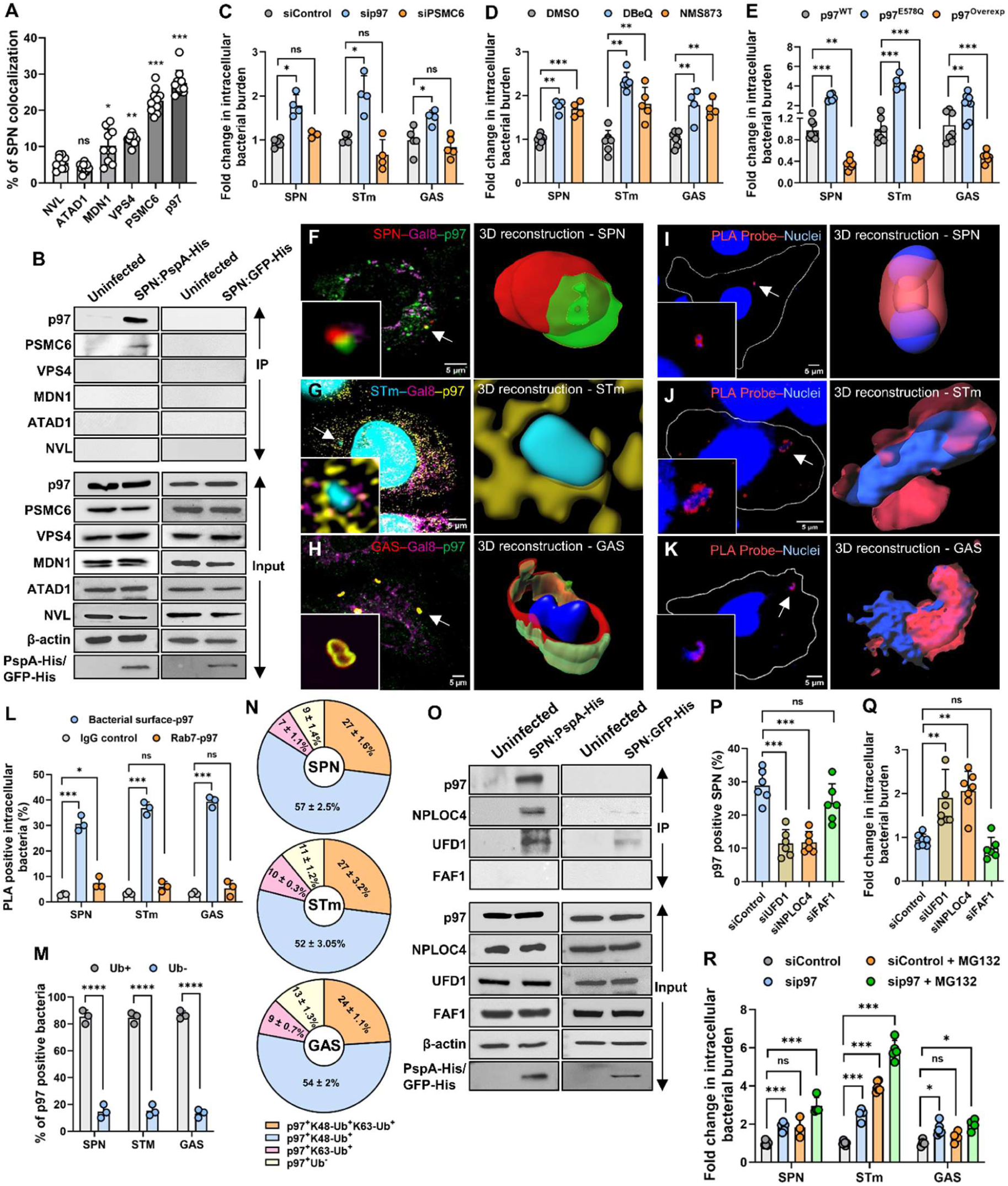
p97 eliminates cytosolic ubiquitinated pathogens. **A.** Quantification of various AAA-ATPases’ (p97, PSMC6, VPS4, MDN1, ATAD1 and NVL) association with intracellular SPN as analysed using immunofluorescence studies. n ≥ 100 events examined over ten independent experiments. Statistical comparisons of bacterial association with different AAA-ATPases were done with NVL as reference. **B.** Immunoprecipitation of pneumococcal surface protein PspA-(His)6 (using anti-His antibody) following infection of A549s with SPN strain Δ*pspA*:pPspA-His demonstrates that the bacterial surface protein interacts majorly with p97, while PSMC6, VPS4, MDN1, NVL and ATAD1 showed minimal or no interaction. SPN expressing GFP-(His)6 was used as a negative control. **C, D, E.** Fold change in intracellular burden of SPN, STm, and GAS, in host cells, either following transfection with sip97 and siPSMC6, normalized to scramblase control **(C)**, or treatment with p97 inhibitors DBeQ and NMS-873, normalized to DMSO (vehicle) **(D)**, or following overexpression of wild type p97 (p97^Overexp^) or p97^E578Q^ (catalytic mutant) **(E)**. A549s are infected with SPN and GAS for 9 h while HeLa cells were infected with STm for 6 h. Fold change in intracellular bacterial burden is depicted as ratio of intracellular bacterial CFU of the test to that of control at specified time points. Data are means ± SD of N = 3-7, independent biological replicates. **F-H.** Structural Illumination Microscopy (SIM) representing the association of p97 (Green or Yellow) with SPN (Red) **(F)**, STm (Cyan) **(G)** and GAS (Red) **(H)** devoid of damaged endosome marker Galectin8 (Magenta). Arrows designate bacteria shown in insets. Galectin8 and p97 were stained with specific antibody and bacteria was stained with either DAPI or surface specific primary Ab, followed by Alexa Fluor 488 or 633 conjugated to secondary Ab. Scale bar, 5 μm. 3D reconstruction of **F**, **G** and **H** using IMARIS are represented on the right side. **I-K.** Confocal micrographs showing association of p97 with surface markers of SPN **(I)**, STm **(J)** and GAS **(K)** as determined by Proximity ligation assay (PLA). Arrows depict bacteria shown in insets. Scale bar, 5 µm. 3D reconstruction of **I**, **J** and **K** using IMARIS are represented on the right side. **L.** Percentage of PLA positive bacteria associating with p97 on their surface (Enolase; LPS; PolyA polysaccharide for SPN, STm and GAS, respectively) vs with endosomes (Rab7). Isotype control Ab served as a negative control. n>50 events examined over three independent experiments. **M.** Percentage of p97-positive bacteria marked with ubiquitin (stained with specific primary antibody). n>50 events examined over three independent experiments. **N.** Percentage of p97-positive bacteria that are decorated with K48-polyUb or K63-polyUb chains. Bacteria were stained with DAPI, p97 was conjugated to GFP, K48-Ub and K63-Ub were stained with respective primary Ab, followed by Alexa Fluor 633 or 555 conjugated to secondary Ab. n>50 events examined over three independent experiments. **O.** Immunoprecipitation of PspA-(His)6 (using anti-His antibody) following infection of A549s with SPN strain Δ*pspA*:pPspA-His, displaying its interaction with NPLOC4 and UFD1, but not with FAF1. SPN expressing GFP-(His)6 was used as a negative control. **P.** Percentage of intracellular SPN marked with p97 following transfection with siNPLOC4, siUFD1 and siFAF1 compared to siControl (scramblase). n>100 events examined over six independent experiments. **Q, R.** Fold change in intracellular burden of SPN upon treatment of A549 cells with siNPLOC4, siUFD1 and siFAF1 compared to siControl treated host cells. **R.** Fold change in intracellular burden of SPN, STm and GAS upon treatment of host cells with either Scramblase (siControl) or sip97 or siControl + MG132 and sip97 + MG132. Fold change in intracellular bacterial burden is depicted as ratio of intracellular bacterial CFU of the test to that of control at specified time points. Data are means ± SD of N = 3-6 independent biological replicates. Statistical significance was assessed by one-way ANOVA followed by Dunnett’s test for **A, C-E, L, N, P-R**. Two-tailed unpaired student’s t-test (nonparametric) was used for **M**. *P < 0.05; **P < 0.01; ***P < 0.005.

Since ATPase activity is key for AAA-ATPase function, cells expressing p97^E578Q^, a catalytically inactive form^26^, not only had higher K48-Ub substrate accumulation, but also had ∼2.3-4-fold higher intracellular bacterial burden (Fig. 1E, S5A). Conversely, overexpression of p97 in host cells lead to improved clearance of the pathogens (∼2-2.8-fold) (Fig. 1E, S5B). Further, polymorphic mutations in p97 (R155H, L198W), alter its activity, leading to multisystem proteinopathy^27^. Cells harbouring these mutant p97 alleles accumulated elevated levels of K48-Ub substrate and had diminished ability to clear intracellular pathogens (∼1.4-2-fold) (Fig. S6). This suggests the possibility that individuals carrying such mutations are at high risk of bacterial infections. Critically, microbial invasion in these mutant cells or cells treated with different pharmacological agents remained similar, nullifying any contribution of infection defect in the perceived pathogenic intracellular burden (Fig. S3I-K, S4I-K, S5C-E, S6C-E).

### VCP/p97 targets cytosolic ubiquitinated pathogens

Since p97 is highly abundant in the cytosol and acts on ubiquitin signals^28^, we speculated that it would target the cytosolic ubiquitinated pathogens. Indeed, ∼50-80% of p97 positive SPN, STm, and GAS were devoid of damaged endosomal marker, Galectin8 or generic membrane marker, FM4-64 (Fig. 1F-H, S7). In addition, 3D-reconstruction of these events allowed visualization of p97’s docking on bacterial surface (Fig. 1F-H). This was further validated by abrogation of p97’s ability to mark a SPN *Δply* mutant lacking the pore forming toxin, pneumolysin, which facilitates bacterial entry to cytosol by puncturing the endosomal membrane (Fig. S2D, E). However, treatment of cells with LLOMe (Lysomotropic agent) enabled decoration of *Δply* with p97 (Fig. S2B, C, E), signifying that cytosolic exposure of the bacteria is required for p97’s recruitment. Moreover, proximity ligation assay using surface markers of different pathogens exhibited significant association of p97 (∼30-40%) with cytosolic bacteria, as compared to the vacuole bound pathogen marked by Rab7 (Fig. 1I-L). Additionally, ∼85% of p97 associated bacteria were found to be ubiquitinated and treatment of cells with PYR41, an E1 inhibitor, reduced microbial association of p97 by ∼40-70% (Fig. 1M, S8A). Further, among the ubiquitin chain topologies, K48-Ub linkage was the primary driver for p97’s microbial targeting (∼52%) (Fig. 1N, S8B-G). Consequently, cells which failed to produce K48-Ub chain type (K48R-Ub) reduced p97’s association with the bacteria (∼2.5-fold), while mutant K63R-Ub had no effect (Fig. S8H). This validates the observed increment in bacterial load in K48R-Ub cell lines^17^. Additionally, knockdown of FBXW7 and ARIH1 (Fig. S3F, H), that synthesize K48-Ub chain on the SPN and STm surface, respectively^12^ ^29^, drastically reduced p97’s association, in contrast to RNF213 downregulated host cells (Fig. S3G) which majorly promotes linear ubiquitin chain^30^ (Fig. S8I, J).

VCP/p97 has weak binding affinity towards its ubiquitinated target and requires specific cofactors for efficient association and functioning^31^. Amongst few K48-Ub targeting cofactors such as NPLOC4, UFD1 and FAF1, significantly high proportion of p97 positive SPN recruited NPLOC4 and UFD1 (∼50-70%), while only ∼30% had FAF1 (Fig. S9A-B). Moreover, the co-association of NPLOC4 and UFD1 (∼65%) on bacterial surface provides evidence of them being the dominating cofactor duo. This trimeric association of p97-NPLOC4-UFD1 on ubiquitinated microbial proteins was further substantiated by their co-immunoprecipitation with PspA (Fig. 1O), where, FAF1 was absent. In addition, knockdown of NPLOC4 and UFD1, but not FAF1 (Fig. S3C-E) significantly reduced bacterial association of p97 and consequently doubled intracellular microbial burden (Fig. 1P-Q). Also, cells expressing p97*^Δ^*^54–241^, having deletion in the hydrophobic pocket containing NPLOC4 and UFD1 binding site, had diminished (∼2-fold) targeting of SPN by p97, which resulted in ∼1.5-fold increase in its intracellular bacterial burden (Fig. S9C-E). Following extraction by p97, K48-polyubiquitinated bacterial surface proteins should presumably be degraded by proteasomes. Indeed, ∼50-90% of p97-positive bacteria also recruited proteasomes (represented by proteasomal subunit β7) (Fig. S10A-D). Reducing the expression of p97 therefore drastically impeded the association of β7 with pathogens (Fig.S10E). Additionally, combination of proteasomal inhibitor (MG132) and p97 depletion elevated intracellular bacterial burden (Fig. 1R), underlining their interplay in guarding the host cytosol.

### VCP/p97 extracts bacterial surface protein and causes deformation in peptidoglycan layer

For the antimicrobial role of p97, we presume that its segregase activity must extract the bacterial surface proteins. Indeed, only ubiquitinated BgaA, immobilized on Sepharose beads was pulled out into the supernatant by p97 (Fig. 2A-B). Additionally, an engineered p97, having a C-terminally fused protease domain of FtsH (FtsHp, from *E. coli*) triggered proteolysis of the extracted BgaA, which was further blocked with protease inhibitor (Fig 2A, C-D). This implies that BgaA is extracted by p97 and released through its C-terminal, where it undergoes degradation by FtsHp.

**Figure 2.**
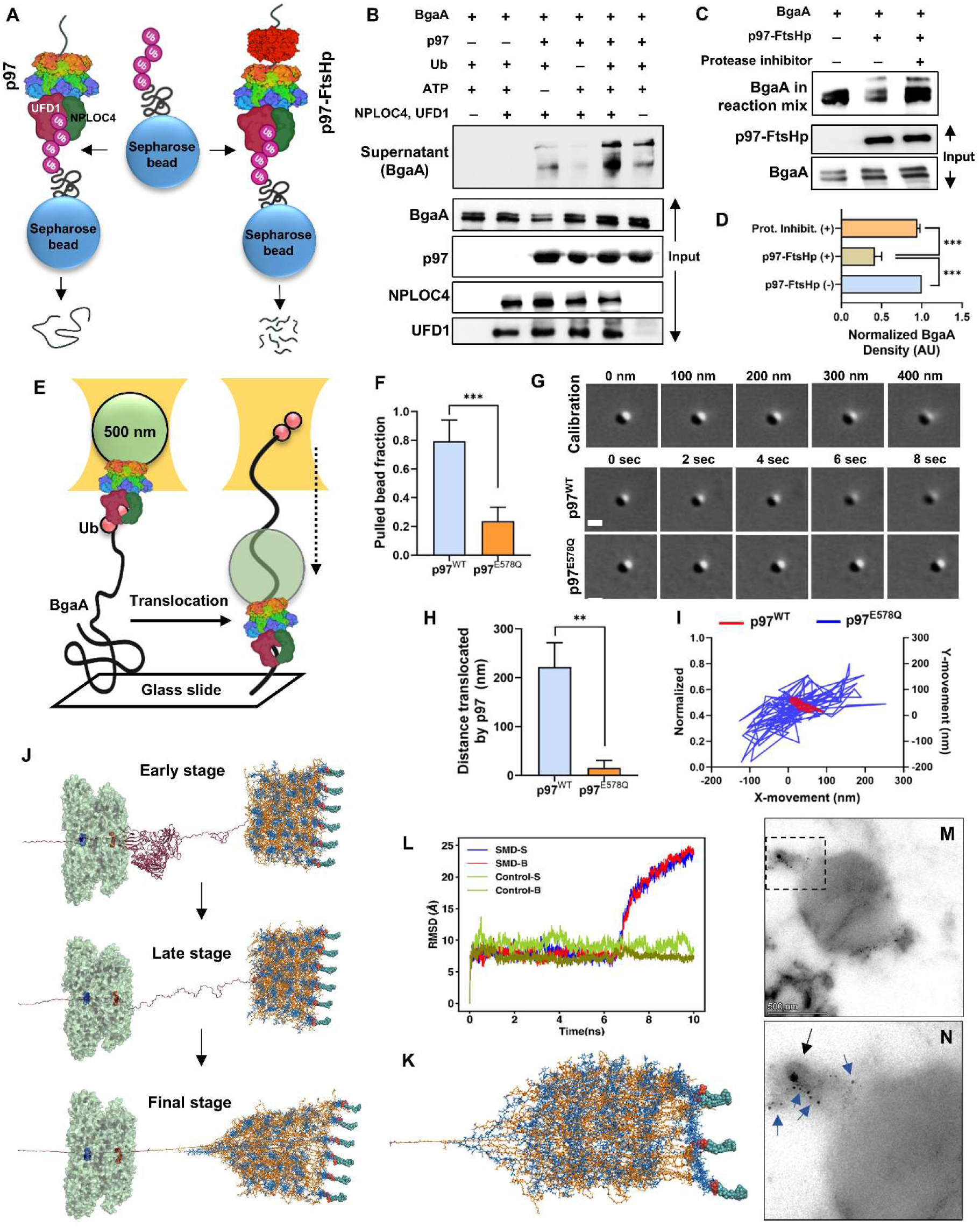
Extraction of bacterial surface protein BgaA by p97 ruptures the cell wall. **A.** Schematic showing the workflow for detecting BgaA upon extraction by p97 or following degradation by p97-FtsHp engineered protein where FtsH protease domain is fused to C-terminal end of p97. **B.** Immunoblot showing the extraction of BgaA bound to sepharose beads by p97 (5 μM) in the mobile/supernatant fraction in presence or absence of Ub, ATP and co-factors NPLOC4 and UFD1. **C.** Immunoblot showcasing the extraction coupled with degradation of BgaA by p97-FtsHp (2 μM) which is impeded upon treatment with protease inhibitor. **D.** Densitometric analysis of BgaA band intensities as observed under different depicted conditions. **E.** Schematic representing optical trap experiment where p97-Strep bound streptavidin beads are allowed to bind to BgaA, immobilized on glass slide, where p97 unfolds and translocate through the length of the substrate (depicted by dashed arrow) in real time. **F.** Fraction of beads conjugated with p97^WT^ or p97^E578Q^ that got pulled towards BgaA, when brought close to the surface using an optical trap within 20 sec. Translocation of n>30 beads over three independent experiments were assessed. **G.** Time-lapse images of representative beads conjugated to either p97^WT^ (top) or p97^E578Q^ (bottom) when binding to BgaA surface to assess translocation. Also shown are images of a control bead physically moved towards the surface of glass slide, represented at every 100 nm step size in Z-axis. Scale bar, 0.5 µm. H. Distance translocated by beads conjugated to p97^WT^ and p97^E578Q^ towards the glass slide surface as calculated from video tracking data (shown in G). Mean Grey scale histogram values were obtained by subtracting intensity of final image (8 sec) by initial mean value (0 sec) in ImageJ and corresponding distances were calculated using the calibration curve (Fig. S11D). I. Representative X-Y trajectory of a bead conjugated with p97^WT^ (red) or p97^E578Q^ (blue) bound to the BgaA surface. Data are means ± SD of N = 8, independent biological replicates. **J.** Representative snapshots from Movie S1, depicting the early, late, and final pulling of the BgaA-p97-peptidoglycan model in molecular dynamic (MD) simulation in combination with steered MD (SMD). In the simulation N-terminal end of surface attached BgaA is pulled through the central pore of the p97 (SMD-S). Similar simulation was also performed with BgaA buried in peptidoglycan layer (SMD-B). **K.** Top view of stretched peptidoglycan following p97 mediated pulling of BgaA as shown in **“J”**. **L.** RMSD time evolution of the sugar units of the peptidoglycan mesh connected to BgaA during SMD-S and SMD-B, in comparison to control simulations. **M.** Immunogold-TEM image showing deposition of p97 on SPN surface. Scale bar, 500 nm. **N.** Zoomed in view of the boxed area in “M” showcasing presence of p97 specific gold particles (blue arrow) and pinching of the peptidoglycan layer (black arrow) from the same area on SPN surface. Statistical significance was assessed by one-way ANOVA followed by Dunnett’s test for **D.** Two-tailed unpaired student’s t-test (nonparametric) was used for F, H. **P < 0.01; ***P < 0.005.

To study the extraction/pulling of BgaA by p97 in real-time, we used the optical trap method (Fig. 2E, S12E, F). To begin with ∼90% of p97 conjugated Streptavidin beads bound to ubiquitinated BgaA immobilized on a hydrophilic coverslip, whereas binding was negligible with non-ubiquitinated BgaA (Fig. S11A). Following binding, individual beads were held in the trap slightly above the coverslip surface. The BgaA coated surface was moved using a piezo at a constant velocity of 300 nm/sec for 1 sec (Fig. 2E). While 80% of p97^WT^ beads bound to the surface and escaped the trap, p97^E578Q^ coated beads bound transiently to the surface, quickly detached and kept falling back to the trap center (Fig. 2F, S11B, C). We video-tracked the binding of beads to the BgaA surface and observed that p97^WT^ beads translocated towards the BgaA, thus shifting below the focal plane towards the coverslip (Fig. 2G). Contrarily, though p97^E578Q^ beads can bind the surface, there is no shift in their focus during binding suggesting no translocation (Fig. 2G). To quantify the displacement of p97 coated beads towards the coverslip, we generated a calibration curve using a strep bead that was stuck to the coverslip and moved downwards in 20 nm steps using a piezo stage (Fig. S11D). Using this plot as the reference, we estimated that p97 beads translocated ∼220 nm toward the coverslip surface, whereas for p97^E578Q^ the displacement was only 10 nm (Fig. 2H). This suggests that during the extraction process, p97 translocates along the length of BgaA while unfolding the protein. If the p97-bound bead moves downwards as p97 generates force to pull in the BgaA substrate, then the bead-to-coverslip tether should reduce, and the bead should wobble less in the lateral direction. Indeed, an X-Y scatter plot showed that p97 beads wobbled significantly less than p97^E578Q^ (Fig 2I, S11E)

We then speculated that the mechanical force generated by p97 during extraction of the substrate, perturbs the cell-wall integrity. To gain insights into this process, molecular dynamic (MD) simulation was employed where either surface attached (SMD-S) or buried (SMD-B) BgaA were pulled through p97’s central channel. This first led to complete unfolding of BgaA followed by deformation of the peptidoglycan mesh (Fig. 2J, K, Movie S1) as evident by the change in Root-mean-squared-deviation (RMSD) of the peptidoglycan layer compared to control simulation (Fig. 2L). Indeed, immunogold-transmission electron microscopy of a SPN inside host cells revealed not only deposition of p97 on bacterial surface, but also pinching off the peptidoglycan layer from SPN membrane (Fig. 2M-N) from the same area.

### VCP/p97 exhibits bacteriolytic activity

We then focused to unambiguously prove the antimicrobial property of this molecular machine by *in-vitro* reconstitution assay using purified p97, p97^E578Q^, NPLOC4 and UFD1 (Fig. S12A-D). Pure p97^WT^ exhibited higher ATPase activity compared to p97^E578Q^ variant and its activity further increased following complex formation with NPLOC4 and UFD1 (Fig. S12G-I), though both variants bind to ubiquitinated proteins (Fig. S12J). Remarkably, viabilities of the *in-vitro* ubiquitinated pathogens (Fig. 3A, S13A) were drastically reduced upon incubation with active p97-NPLOC4-UFD1 complex. However, non-ubiquitinated or deubiquitinated (treatment with OTUB1) or p97^E578Q^-NPLOC4-UFD1 treated bacteria escaped p97 mediated eradication (Fig. 3B). Additionally, CFU count of SPN, ubiquitinated with purified E1, E2 and Rbx1-Cul1-SKP1-FBXW7 (E3 ligase), showed significant reduction following treatment with p97, which was robbed off upon treatment with deubiquitinase (OTUB1) (Fig. S13B). We also determined the kinetics as well as half inhibitory concentration (IC50) of p97 required in this process (Fig. 3C-D). To prove whether extraction of ubiquitinated proteins by p97 punctures the bacterial membrane, leading to their death, we analysed permeability of Sytox orange (membrane impermeable dye). Indeed, Sytox intake was primarily on p97 positive pathogens (Fig. 3E). Additionally, we found significant increase (∼4-fold) in Sytox positive bacterial population when incubated with active p97 compared to that of denatured or p97^E578Q^ complex (Fig. 3F-H, S14). Further, visualization of bacterial surface morphology by scanning electron microscopy showcased profound rupture or membrane blebbing of all three pathogens when subjected to active p97 treatment (Fig. 3I). Such large-scale disintegration of the bacterial membrane may lead to release of the internal contents, including DNA. Indeed, active p97 triggered excessive leaching of bacterial DNA (∼2-10-fold) while catalytically inactive or denatured complex showed negligible release (Fig. 3J-L). These confirmed specific targeting of ubiquitinated pathogens by p97 and the indispensable role of D2 domain associated ATPase activity in generating mechanical force for exhibition of bactericidal activity.

**Figure 3.**
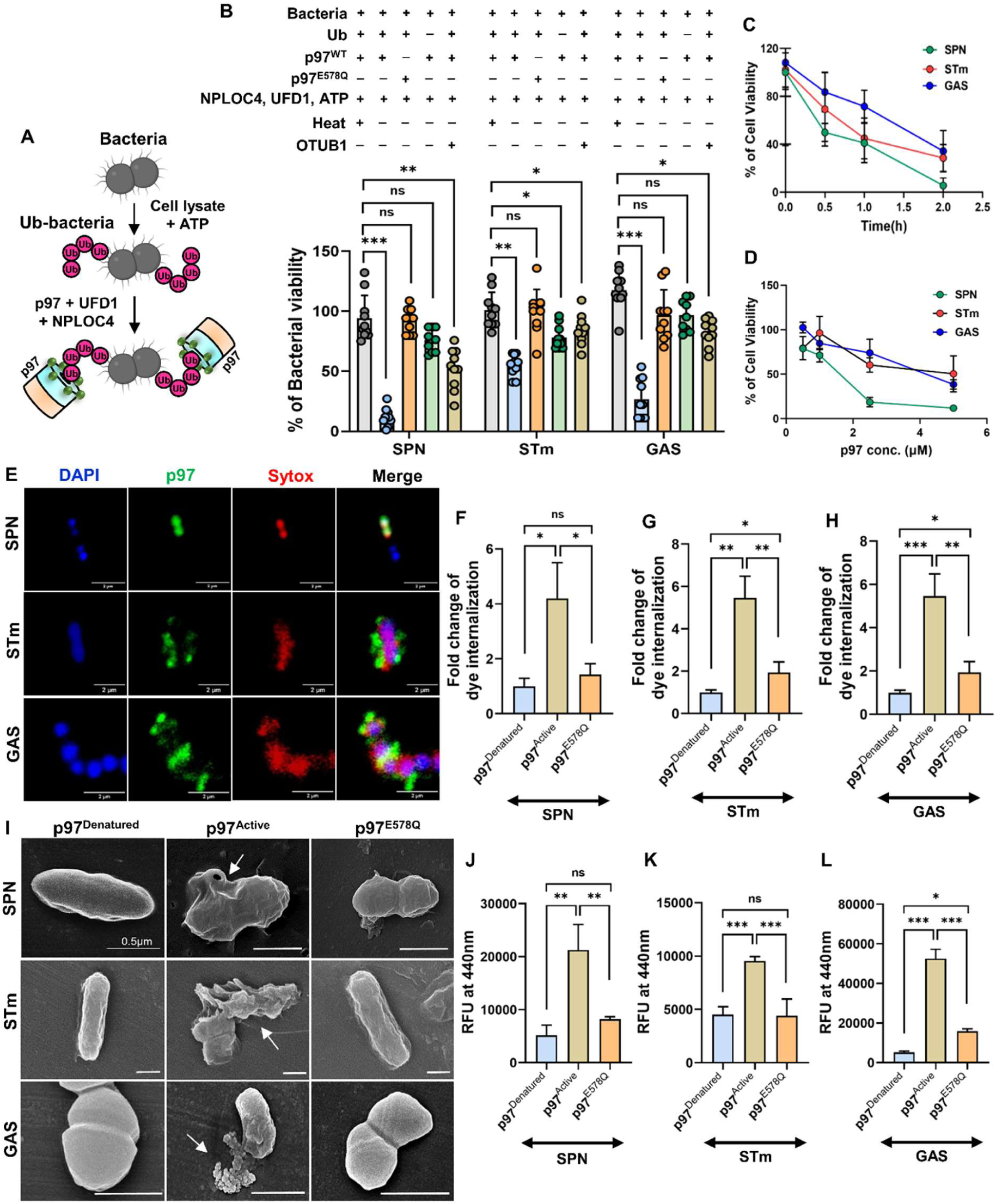
p97 showcases bacteriolytic ability *in-vitro*. **A.** Schematic representation showing the workflow for *in-vitro* reconstitution of bacterial ubiquitination followed targeting by p97. **B.** Percentage of bacterial cell viability upon treatment with denatured (p97^Denatured^) and active p97 complex (p97^Active^) along with catalytic mutant, p97^E578Q^. SPN, STm and GAS with or without *in-vitro* ubiquitination were incubated with p97^WT^, p97^Denatured^ or p97^E578Q^ (5 µM) in presence of co-factors NPLOC4 and UFD1 (∼1 µM each) for 2 h. Bacterial numbers in the reaction mix was estimated by spread plating. Ubiquitinated bacteria were also treated with deubiquitinase, OTUB1, to assess effect of ubiquitination on bacterial targeting by p97. Data are means ± SD of N = 10 independent biological replicates. Percentage of bacterial viability was enumerated by: (final CFU/initial CFU) × 100%. **C.** Killing kinetics of p97 complex (5 µM) against various bacteria. Data are means ± SD of N = 3 independent biological replicates. **D.** Viability of SPN, STm and GAS following incubation with p97 and co-factors for 2 h over a concentration range (0.5 - 5 µM) to determine the half maximal inhibitory concentration (IC50). Data are means ± SD of N = 3 independent biological replicates. **E.** Confocal micrographs showing the internalization of a membrane-impermeable dye, Sytox orange (Red), in bacteria (Blue) upon association with p97 (Green). Bacterial DNA was stained with Hoechst (Blue). Scale bar, 2 µm. **F-H.** Flow cytometry analysis showing the extent of Sytox internalization in a population of SPN **(F)**, STm **(G)** and GAS **(H)** cells following incubation with active p97 complex compared to denatured control and p97^E578Q^. Corresponding scatter plots depicting percentages of Sytox positive bacteria are shown in Fig. **S14**. n>40000 events examined over three independent experiments. **I.** Scanning electron microscopy displaying surface disintegration of bacteria upon incubation with active p97 complex, compared to denatured control and p97^E578Q^. Scale bar, 0.5 μm. **J-L.** DNA release from SPN **(J)**, STm **(K)** and GAS **(L)** upon treatment with active p97 complex compared to denatured control and p97^E578Q^. Ubiquitinated bacteria were stained with Hoechst 33342 (5 µg/ml) and treated with p97. Released DNA were estimated by assessing fluorescence of the supernatant. Data are means ± SD of N = 3 independent biological replicates. All experiments were performed with purified proteins. Heat denatured protein complex was used as a negative control. Statistical significance was assessed by one-way ANOVA followed by Dunnett’s test. ns, non-significant; *P < 0.05; **P < 0.01; ***P < 0.005.

### VCP/p97 protects the host from severe bacterial infection

We then sought to evaluate p97’s ability as a guarding factor using an established pneumococcus infection induced *in-vivo* sepsis model (Fig. 4A)^32^. Inhibition of p97 by NMS-873 led to increased accumulation of K48-Ub substrates (Fig. S15A) and significantly higher mortality of host than vehicle cohort (p=0.0072, log-rank test). While all the p97 inhibited animals succumbed to infection within 38 h, >40% of vehicle treated mice survived during the course of the experiment (median survival time 18 h and 30 h, respectively) (Fig. 4B). Moreover, bacterial burden in blood and spleen of the p97 inhibited mice at the time of death was ∼10-15-fold higher than the vehicle treated animals (Fig. 4C, D). In sepsis, following initial clearance of the pathogen from the bloodstream, a minor subset of SPN survives and clonally expands in few splenic macrophages, eventually re-seeding into the blood, causing fatal infection^32^. We speculated that p97 could impede this process. Indeed, both in spleen and blood, SPN burden kept increasing with time (∼10-12-fold) in the NMS-873 treated cohort at 12 h.p.i (Fig. 4E, F). Further, immunohistochemistry studies revealed that splenic burden of SPN in p97 inhibited animals was ∼4-fold higher and reached as high as 10 bacteria per foci (∼300 μm^2^), in contrast to majorly single SPN in case of vehicle cohort (Fig. 4G-K). This indicates rapid clonal expansion of SPN in absence of immune surveillance by p97. Further, increased level of proinflammatory cytokines IL-6 and IL-1β was found in blood and spleen in the p97 inhibited mice (Fig. 4L, M). Additionally, treatment of host with LPS in presence and absence of NMS-873 did not alter the cytokine levels in serum and spleen (Fig. S15B-C), ruling out the possibility of general immune suppression in host due to p97’s inhibition. Thus, inhibition of p97 exacerbates pathogen induced septicemia like conditions and demonstrates its utmost importance in host protection against microbial pathogenesis.

**Figure 4.**
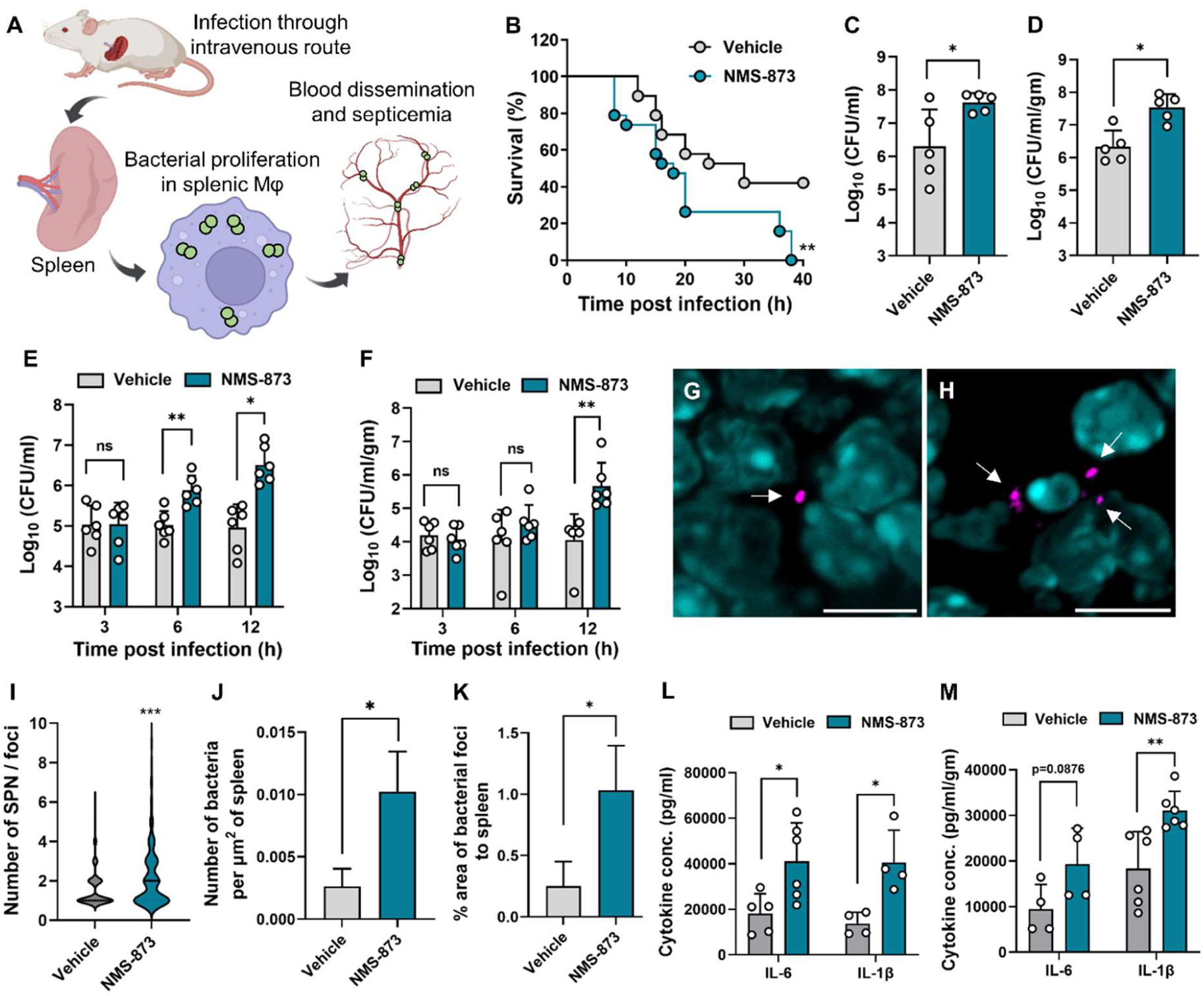
p97 protects mice from bacterial sepsis. **A.** Schematic describing mouse model of septicemia following pneumococcal infection. **B.** Kaplan-Meier survival curve of mice after i.v. infection with SPN (10^6^ CFU) and treatment with either p97 inhibitor NMS-873 (0.02 mg/kg) or vehicle. n=19 mice per group. **C-D.** Bacterial burden at the time of death in blood **(C)** and spleen **(D)**. n=5 mice per group. **E-F.** Comparison of bacterial burden in blood **(E)** and spleen **(F)** of mice treated with either NMS-873 or vehicle at regular intervals during the first 12 h.p.i with SPN. **G-H.** Representative immunohistochemistry images showing clustering of SPN in spleen of NMS-873 treated mice **(H)** compared to control cohort **(G)**. Arrows depict SPN (magenta, stained with Anti-enolase Ab) in spleen section stained with DAPI. Scale bar, 10 µm. **I.** Number of SPN per foci, in spleen of animals treated with either NMS-873 or vehicle cohort. **J-K.** Number of bacteria per area (μm^2^) of spleen **(J)** and percentage of area occupied by SPN **(K)** in spleen slices from animals treated with NMS-873 compared to vehicle. n=3. **L-M.** Cytokine levels in blood **(L)** and spleen **(M)** at 12 h.p.i. Experiments were repeated twice and data is collectively represented. Statistical analysis was assessed using two-tailed unpaired student’s t-test (nonparametric). ns, non-significant; *P < 0.05; **P < 0.01; ***P < 0.005.

## Discussion

Host autonomous immunity has evolved as a consequence of pathogenic challenges. The intent to survive in adversity has driven the development of a gauntlet of efficient immune surveillance systems to detect and defend against all foreign threats. Often invasive pathogens infiltrate host cells and escape lysosomal fusion by modifying the vacuoles harboring them. Accidental or purposeful rupture of such phagosomes leads to release of pathogens into the cytosol. This allows access to nutrient-rich cytosolic environment, which not only could promote microbial growth, but also encourage evasion from vacuolar death traps. However, upon escape into the host cytosol, interferon-induced GBPs perforate pathogen’s membrane, inducing pyroptosis^6, 9, 13, 14, 33, 34^. Although the role of GBPs is versatile, its selectiveness to the pathogen’s origin, especially towards Gram-negative bacteria, projects an incomplete picture about the diversity and nonspecific nature of the cytoplasmic innate immune mechanisms. This underlines the presence of an unbiased immune sentinel that protects the metazoans against broad-spectrum intruders. Our work unveils a unique bacteriolytic activity of an AAA-ATPase, p97, which acts against diverse cytosol dwelling microbes. By generating mechanical force, these nanomachines extract ubiquitinated substrates common to both Gram-positive and Gram-negative bacteria, leading to their lysis. This unusual generic mechanism employed by the host limits resource diversification and sheds light on the fundamental role of AAA-ATPase family in thwarting a variety of microbial infections. Recently, p97 has been implicated in targeting of parasite containing vacuoles and antibody-dependent intracellular neutralization of viral particles^35, 36^. However, we observed direct targeting of the microbial surface by p97, which is driven by K48-polyUb conjugation to bacterial surface proteins. The ability of cellular ubiquitination machinery to label a variety of microbes with ubiquitin, drives this impartial bacteriolytic capacity of p97.

Invasive pathogens often establish systemic infections by replicating in splenic macrophages from where they re-seed into the circulatory system to overwhelm the host^32, 37^. Interestingly, our results emphasize a key role of p97 in impeding bacterial replication in the spleen and subsequent dissemination into the vasculature. This highlights the potential role of cytosolic guarding factors in averting microbial proliferation in general antigen presenting cells or other cell types which might be absent in immunocompromised sites. We further postulated that mutations in p97 leading to inept functioning might compromise microbial clearance. In fact, patients carrying such mutations suffer from inclusion body myopathy with early onset of Paget disease and frontotemporal dementia (IBMPFD) and are notably linked to disruption of various homeostasis processes^38^. Remarkably, we observed a striking relation between host cells expressing such hotspot mutations of p97 with profound inability to clear pathogens, leading to escalated infection susceptibility. This correlation between polymorphism in individuals and their vulnerability to infections has also been noted in patients afflicted with Parkinson’s and chronic lymphocytic leukemia (CLL)^12, 39^. Collectively, these serve as a strong reminder that genetic makeup of individuals can influence the on-going battle against pathogenic adversaries and have a cardinal impact on innate defence.

In conclusion, we unravel an ancient yet powerful class of host immune factor that plays a decisive role in clearance of diverse pathogenic bacteria. Exploitation of such broad-spectrum antimicrobial mechanism could tip the balance in favour of the host against ever evolving bacterial and non-bacterial foes.

## Acknowledgement

We acknowledge the "Bio-safety Level 2 Facility" and “Confocal Microscopy Facility” at IIT Bombay. We also acknowledge Electron microscopy facility at CSIR-CCMB, Hyderabad, India, and Animal facility at IISc, Bengaluru, India.

## Author contributions

### Conceptualization

AB

### Methodology

SG, SR, NB, UKD, PS, DH, SM, MS, HA

### Validation

SG, SR, NB, UKD, PS, DH, SM, MS

### Formal Analysis

SG, SR, PS, DH, SRa, SM, MS, JM, RM, AB

### Investigation

SG, SR, NB, UKD, PS, DH, SRa, SM, MS, SS, RSR, HA

### Resources

JM, DC, RM, AB

### Data Curation

SG, SR, PS, JM, RM, AB

### Writing – Original Draft Preparation

SG, AB

### Writing – Review & Editing Preparation

SG, NB, SR, SS, AB

### Visualization

SG, SR, NB

### Supervision

JM, DC, RM, AB

### Project Administration

AB

### Funding Acquisition

AB

## Supplementary Materials

### Materials and Methods

Figures S1-S15

Tables S1-S3

Movie S1

References

