## Supplemental information for "A host AAA-ATPase exhibits bacteriolytic activity for clearance of microbial infection"

**This file contains:**

Materials & Methods
Figures S1 to S15
Table S1 to S3
Movie S1

### Materials and Methods

#### Antibodies and Reagents

Antibodies against VCP/p97 (MA3-004), NPLOC4 (PA554749), PSMB7 (PA5-111404), PSMC6 (PA5-60140), β-Actin (MA5-15739), Group A *Streptococcus* (PA1-73059), MDN1 (PA5-56225), FAF1 (PA5-121137) were purchased from ThermoFisher Scientific. Antibodies specific for UFD1 (Santa Cruz Biotechnology, sc-377265), ARIH1 (Santa Cruz Biotechnology, sc-514551), K48-Ub linkage (MERCK, 05-1307), Phosphothreonine (Cell Signaling, Technology 9381S), K63-Ub linkage (Enzo, BMLBML-PW0600-025), *Salmonella* O antisera (Difco, 225341), Anti-Enolase (Prof. S. Hammerschmidt, University of Greifswald, Germany), VPS4 (MERCK MABC83), NVL (Sigma-Aldrich HPA028219), RNF213 (Sigma-Aldrich, HPA003347), ATAD1 (Sigma-Aldrich, HPA037569) were used in the study. HRP tagged anti-rabbit (Biolegend 406401), HRP tagged anti-mouse (Biolegend, 405306), anti-rabbit Alexa Fluor 555 (Invitrogen, A31572), Anti-rabbit Alexa Fluor 488 (Invitrogen, A27206), anti-mouse Alexa Fluor (Invitrogen, A31570), anti-goat Alexa Fluor 633 (Invitrogen, A21082), anti-mouse Alexa Fluor 488 (Invitrogen, A21202), Biotin conjugated anti-mouse IgaA (Life Tech., M31115), were used as secondary antibody.
Pharmacological inhibitors and small molecule reagents such as DBeQ (SML0031), NMS-873 (SML1128), PYR41 (N2915), and LLOMe (L1002) were purchased from MERCK. Dyes such as FM4™-64 (T3166) and SYTOX™ Orange Dead Cell Stain for flow cytometry (S34861) were purchased from ThermoFisher Scientific.

#### Cell culture and Transfection

Human lung alveolar carcinoma (type II pneumocyte) cell line A549 (ATCC No. CCL-185) and cervical adenocarcinoma cell line HeLa (ATCC No. CRM-CCL-2) were grown in DMEM (HiMedia, AT007) which was supplemented with 10% FBS (GIBCO, 38220090) at 5% $CO_2$ at 37°C. Transfections were performed using Lipofectamine 3000 (ThermoFisher Scientific, L3000001) according to the manufacturer's protocol, and selections were done in the presence of 2 µg/ml blasticidin hydrochloride (HiMedia, TC421).

#### Bacterial Strains and Growth Conditions

All bacterial strains used in the study are listed in Table S1. *E. coli* (DH5α, BL21(DE3), Rosetta (DE3)) cultures and *Salmonella enterica* Serovar Typhimurium (*STm*) (ATCC 14028) were grown in Luria-Bertani broth (LB) at 37°C under shaking conditions (200 rpm). *Streptococcus pneumoniae (SPN)* strains R6 (serotype 2, unencapsulated) and

D39 (serotype 2, capsulated) (provided by Prof. TJ Mitchell, Univ. of Birmingham, UK) and *Streptococcus pyogenes (GAS)* strain JRS4 (obtained from Prof. Victor Nizet, Univ. of California-San Diego, USA) were grown at 37°C with 5% $CO_2$ in Todd-Hewitt broth (THB) supplemented with 1.5% yeast extract. Following antibiotics were used when required: ampicillin (100 µg/ml, for *E. coli*), kanamycin (50 µg/ml for *E. coli* and 200 µg/ml for SPN), chloramphenicol (20 µg/ml for *E. coli* and 4.5 µg/ml for SPN) and spectinomycin (100 µg/ml for SPN).

**Gentamicin protection assay**

*SPN, STm and GAS* grown till $OD_{600nm}$ 0.4 were used to infect monolayers of A549s (with SPN at MOI ~ 10 and GAS at MOI ~ 2) or HeLa (with STm at MOI ~ 50) for 1 h. After washing, extracellular bacteria were killed using antibiotics (10 µg/ml penicillin and 400 µg/ml gentamicin for SPN or 100 µg/ml gentamicin for STm and GAS) for 2 h. Following washes with 1X PBS, cells were trypsinized (0.025% trypsin EDTA), lysed (0.025% Triton X-100), and serial dilutions of the lysates were plated on Brain Heart Infusion (BHI) agar plates to enumerate bacterial colonies. Bacterial invasion efficiency was calculated as (recovered CFU/initial inoculum CFU) × 100%. To calculate the intracellular survival of the bacteria, infected cell lysates at different time points were spread-plated, enumerated, and represented as percent survival at indicated time points relative to 0 h (post-antibiotic treatment). The ratio of survival efficiencies of the test to that of control was calculated and represented as fold change in intracellular bacterial burden.

**Fluorescence Microscopy**

A549 and Hela cells grown on glass coverslips were infected with *SPN (*MOI ~ 25)*,* *GAS* (MOI ~ 2), and *STm* (MOI ~ 50). At desired time points post-infection, cells were washed with 1X PBS, and fixed with ice-chilled Methanol for 10 min at -20°C. Further, 3% BSA was used for blocking for 2 h at RT. Cells were then treated with the appropriate primary antibody in 1% BSA overnight at 4ºC, followed by incubation with a suitable secondary antibody in 1% BSA for 1 h at RT. Finally, coverslips were washed with PBS and mounted on glass slides along with VectaShield® with or without DAPI (Vector Laboratories) for visualization using a Laser Scanning Confocal microscope (Carl Zeiss LSM 780) under 40X or 63X oil objective or using Nikon AXR microscope for Instant-Structured Illumination Microscopy (iSIM) at a 60X or 100X oil objective. The images were acquired after optical sectioning and then processed using Zen lite software (Version 5.0.).

**Proximity Ligation Assay**

For *ex-vivo* proximity ligation, SPN infected A549s were fixed with chilled methanol followed by further processing with Duolink® In Situ Orange Starter Kit Mouse/Rabbit (MERCK, DUO92102-IKT) according to the manufacturer's protocol. Fixed cells were imaged as mentioned above and the colocalization events were quantified using Zen Blue 3.7.

**Transmission electron microscopy**

Following infection of A549 and Hela cells with SPN, GAS and STm, respectively, cells were washed with 0.1 M sodium cacodylate buffer, fixed with 2% paraformaldehyde and 0.5% glutaraldehyde in 0.1 M phosphate buffer, pH 7.4 at room temperature for 2 h. Cells were then washed thoroughly, collected through a cell scraper, and pelleted at 2000 rpm for 10 min. Gradually dehydration was performed with gradient of ethanol (50%, 70%, 90%, and 100% for 15 min), followed by infiltration in ethanol and LR white resin mixtures at 1:1 and embedding in pure LR white resin (Electron Microscopy Sciences, 14300) by polymerization at 65°C for 24 to 48 h. Ultrathin sections (70 nm) were cut using a diamond knife on a Leica EM UC7 Ultramicrotome. The grids were transferred onto drops of the matching Aurion-blocking solution for 15 min and washed with PBS followed by incubation with p97 primary and 25 nm gold conjugated secondary antibody (both at 1:50 dliution) for 1 h at room temperature. The grids were contrasted with uranyl acetate for 15 min and sections were viewed using a transmission electron microscope (Talos L120 C, Thermo Fisher Scientific) at 120 kV.

**Vector constructs**

For transfection, EGFP-p97 gene was amplified from pEGFP-p97 (Addgene #85670) and cloned into pMRX vector (Gifted by T Yoshimori, Osaka University, Japan). Further site directed mutagenesis was performed to create various p97 variants in the pMRX background mentioned in table S2, S3.
For protein purification, p97 gene was amplified from pMRX-EGFP-p97 and cloned into pET28a vector along with $p97^{E578Q}$ variant which was further transformed into *E. coli* BL21(DE3). Protease domain of FtsH (FtsHp, 401-664 a.a.) from *E. coli* (DH5α) was translationally fused at the c-terminal region of p97 in pET28a. We added 6X-His at the C-terminal end of NPLOC4 and cloned into pET41b followed by transformation into Rosetta (DE3) (Table S2, S3).

**Protein expression and purification**

*E. coli* Bl21(DE3) strain carrying p97 or p97 variant (pAB713 and pAB730, pAB1003 respectively) or *E. coli* Rosetta(DE3) harboring NPLOC4 and UFD1 (pAB720 and

Addgene #117107, respectively) were cultured in LB as described above. The overnight cultures were re-suspended in fresh LB (1:10) and were incubated at 37°C. At $OD_{600nm}$ ~ 0.8, isopropyl-1-thiogalactopyranoside (IPTG) (200 µM for p97 and its variants, 1 mM for NPLOC4 and UFD1, respectively) was added to the cultures, followed by further incubation for 5 h at 37°C. For NPLOC4 expression, post IPTG addition cultures were further incubated at 18°C for 16 h. The bacterial suspensions were pelleted down at 4000 rpm for 15 min and re-suspended in lysis buffer (50 mM Tris-Cl, 300 mM NaCl, 5 mM β-mercaptoethanol, 1 mM PMSF, 5% glycerol, pH.8) and lysed by sonication. The sonicated lysates were centrifuged at 15000 rpm for 30 min and the collected supernatants were loaded onto the equilibrated Ni-NTA column. Columns were washed with buffer (50 mM Tris-Cl, 300 mM NaCl, 5 mM β-mercaptoethanol, 5% Glycerol) containing 30 µM imidazole and bound proteins were with 250 µM imidazole. The eluted fractions were pulled, concentrated with Amicon ultrafiltration unit (10 kDa) and buffer exchanged in 200 mM HEPES, 1 M KCl, 100 mM $MgCl_2$, pH 7.4, using PD-10 columns (GE Healthcare).

**Western blotting**

Mammalian cells were lysed in ice-cold RIPA buffer (50 mM Tris-Cl, pH 7.89, 150 mM NaCl, 1% Triton X-100, 0.5% Sodium deoxycholate, 1% SDS) containing protease inhibitor cocktail (Promega), Sodium fluoride (10 mM) and EDTA (5 mM). Protein was estimated using Bradford reagent (ThermoFischer Scientific), separated on 6-12% SDS-PAGE gels after adding the Laemmli SDS loading buffer and transferred to activated PVDF or nitrocellulose membrane. Finally, the membrane was blocked with 5% skimmed milk and probed with appropriate primary and secondary HRP-conjugated antibodies. Visualization of the blots were done using ECL substrate (Bio-Rad) using a Chemidoc (BioRad).

**Immunoprecipitation**

A549 cells were infected with SPN strain Δ*pspA*:pPspA-His, expressing PspA-$(His)_6$ for 9 h followed by cellular lysis using RIPA buffer. Equal amounts of lysate of infected and non-infected cells were incubated with 5 µg of anti-$(His)_6$ antibody overnight at 4˚C. 40 µl of Protein G Sepharose® was added to the lysate and incubated for 3 h at 4˚C. The beads were then centrifuged at 7000 rpm for 3 min and washed thoroughly. Finally, the sample was separated in SDS-PAGE after addition of Laemmli SDS loading buffer and transferred onto the PVDF membrane followed by probing with specific antibodies for visualization under Chemidoc (BioRad).

**ATPase assay**

The ATPase activity of p97 was measured using the Malachite Green Phosphate assay kit (Sigma, MAK307) according to the manufacturer's protocol in a reaction buffer containing 200 mM HEPES, 1 M KCl, 100 mM $MgCl_2$, and 10 µM ATP, pH 7.4. Absorbance was measured at 620 nm and normalized to 0 h and the turnover of free phosphate was calculated.

***In-vitro* ubiquitination assay**

*In-vitro* ubiquitination was performed by incubating bacteria with host cell lysate. A549 cell lysate was prepared by scraping the cells from T75 flasks at 4°C, followed by sonication (amplitude 40 mA; 2-sec on followed by 2-sec off) in reaction storage buffer (200 mM HEPES, 1 M KCl, 100 mM $MgCl_2$, and 5% Glycerol, pH 7.4). The supernatant was collected from the sonicated product by centrifuging at 15000 rpm for 45 min, followed by flash freezing (in liquid nitrogen).
100 µl of mid-exponentially grown bacteria (approximately $10^7$ cells/ml) was centrifuged, and the pellet was incubated with A549 cell lysate (2 mg/ml total protein) along with 1 mM ATP for 1-2 h at room temperature. The bacteria were washed, fixed with PFA, followed immunostaining with ubiquitin and bacterial surface marker specific antibody. Consequently, Alexa Fluor 488 or 555 conjugated to secondary Ab was added and the sample was mounted using Vecta-shield with DAPI (Vector Laboratories) and visualized under a Laser Scanning Confocal microscope (Carl Zeiss LSM 780).
Alternatively, purified 6x-His-ubiquitin (350 µM, UBPBio, E1830), UBE1 (400 nM, UBPBio, B1100), UbE2C (3 µM, UBPBio, C1300), Rbx1-Skp1-Cul1-Fbxw7 complex (500 nM, Sigma, 23-030), ATP (2 mM) was incubated with actively growing SPN for 2 h at 37°C in reaction buffer consisting of 200 mM HEPES, 1 M KCl, 100 mM $MgCl_2$, and 5% Glycerol, pH 7.4.

***In-vitro* extraction and degradation assay**

For *in-vitro* extraction assay, purified BgaA-T-$(His)_6$ (1-1049 a.a. of BgaA) was immobilized on Sepharose beads and ubiquitinated by treatment with cell lysate (2 mg/ml total protein) along with 1 mM ATP in the reaction buffer at room temperature for 1 h. Following this, p97 (5 µM) along with NPLOC4 and UFD1 (~1 µM) was added with 1 mM ATP and incubated for 2 h. The beads were centrifuged at 15000 rpm for 5 min and the supernatant fraction was subjected to immunoblotting with anti-$(His)_6$ antibody.
For degradation assay, p97-FtsHp (2 µM) (protease domain) along with NPLOC4 and UFD1 (ratio 6:1:1) was incubated with sepharose beads coated with ubiquitinated BgaA-T for 2 h at 37°C in presence of 1 mM ATP. The reaction was then boiled and

separated in Laemmli SDS loading buffer followed by transfer into PVDF membrane and visualized by probing with anti-$(His)_6$ antibody.

***In-vitro* killing assay**

Purified p97 (5 µM), NPLOC4 (~1 µM), and UFD1 (~1 µM) were mixed in a 6:1:1 ratio and incubated on ice for 3 h in a buffer containing 200 mM HEPES, 1 M KCl, 100 mM $MgCl_2$, 5% glycerol and 1 mM ATP (reaction buffer). Ubiquitinated bacteria were incubated for 2 h with the p97 complex at 37°C, followed by colony enumeration on BHI or LA plates and the percentage of bacterial cell viability was calculated by: (final CFU/initial CFU) × 100%.

**Scanning electron microscopy**

Ubiquitinated *SPN, STm and GAS* were incubated with p97-UFD1-NPLOC4 complex for 1 h. Bacterial cells were then fixed with 2.5% glutaraldehyde for 3 h at room temperature, followed by repeated washing with distilled water. The samples were serially dehydrated with 10%, 30%, 50%, 70%, and 100% ethanol, spotted and air dried on an aluminium tray followed by imaging via (Field emission gun) FEG-SEM (JSM 7600F).

***In-vitro* DNA release**

Ubiquitinated bacteria were incubated with Hoechst 33342 (5 µg/ml) (ThermoFisher Scientific) for 10 min to stain the DNA. Stained bacterial suspension was mixed with VCP/p97 complex and incubated for 2 h at 37°C. Post incubation, bacterial suspensions were centrifuged and relative fluorescence unit (RFU) of the supernatants collected were determined by a spectrofluorimeter with an excitation wavelength of 360 nm and emission at 440 nm.

**Flow Cytometry**

After treatment of ubiquitinated bacteria ($10^7$ CFU) with p97 complex, cells were washed with 1X PBS, followed by incubation with SYTOX™ Orange Dead Cell Stain (ThermoFisher Scientific) for 20 min according to the manufacturer's protocol. Flow cytometry was performed using a BD FACS Aria-Fusion flow cytometer. At least 40000 events were collected to detect Sytox-positive bacteria with an excitation wavelength of 561 nm. For analysis, FlowJo (Version 9.3) was used.

### RNA interference

For siRNA-mediated gene knockdown, ON-TARGET PLUS SMARTpool specific for VCP/p97 (L-008727-00-0010, 10 and 15 pmol), UFD1 (L-017918-00-0005, 10 pmol), NPLOC4 (L-020796-01-0005, 15 pmol), FAF1 (L-009106-00-0005, 15 pmol), RNF213 (L-023324-00-0005, 15 pmol), ARIH1 (019984-00-0005, 15 pmol), FBXW7 (L-004246-00-0005, 10 pmol) and PSMC6 (L-009570-01-0005, 15 pmol) were purchased from Dharmacon. A549 and Hela cells were plated in a 24-well plate followed by appropriate siRNA transfection using Lipofectamine RNAiMAX (ThermoFischer Scientific) according to the manufacturer's protocol.

### Molecular Dynamics Simulation

Molecular dynamics (MD) simulation was used to model the disruption of the structural integrity of bacterial peptidoglycan upon extraction of the ubiquitinated bacterial surface protein, BgaA, through the pore of the host p97 hexamer. The structure of the human p97 hexamer was derived from protein database (PDB ID: 7BP8) and structure of truncated version (97-1036 a.a.) of BgaA, was predicted using AlphaFold2[40]. Adopting an all-atom model of *Staphylococcus aureus* peptidoglycan, we built a 3D peptidoglycan mesh consisting of 6 x 5 chains of peptidoglycan crosslinked by pentaglycine chains. We performed implicit solvent simulations of the system using the generalized Born implicit solvent model as implemented in NAMD 2.14 and performed at a timestep of 1 fs. The CHARMM36m force field is used for the proteins BgaA and p97. An $\alpha$ cutoff, corresponding to the Born radius of the atoms is set as 12 Å. The nonbonded Lennard Jones interactions are cut off at a distance of 14 Å with a switching function at 13 Å. The system temperature is maintained at 310 K using a Langevin thermostat. In order to simulate the process of extraction of BgaA through the pore of p97 hexamer, we harnessed the technique of steered MD simulations, which is an enhanced sampling technique for sampling rare events. SMD simulations were performed with the bactoprenol groups fixed and the C$\alpha$ atom of K97 in BgaA subjected to pulling at constant velocity. The simulation time scale was 1 ns and 10-12 ns using pulling velocities of 0.005 Å/timestep (equivalent to 5 Å/ps) and 0.0005 Å/timestep (equivalent to 0.5 Å/ps). Visualization and analyses of the simulation were performed using VMD and TCL scripting interface.

### Optical trap

1mg/ml BgaA protein was added to a flow chamber made with an acid washed coverslip. The protein was allowed to be adsorbed on the coverslip for 3 h at RT. The unbound protein was washed off and BgaA was ubiquitinated by adding 4 µg/µl of A549 cell lysate for 1 h. After washing, the coverslip was incubated with 5% casein in reaction buffer (200 mM HEPES, 1 M KCl, 100 mM $MgCl_2$, 5% glycerol and 1 mM

ATP). 10 µl of Streptavidin beads (Spherotech SVP-05-10) were washed and incubated with 5 µg of p97 or p97$^{E578Q}$ protein on ice for 15 min. To remove unbound p97, bead protein mix was centrifuged at 10,000 rpm for 5 min. The pellet was resuspended in 500 µl of reaction buffer and bath sonicated for 15 min. Before flowing the beads on the chamber, they were vortexed vigorously to avoid clumping. For the assay p97 (5 µM) conjugated beads were mixed with cofactors (UFD1 and NPLOC4, 1 µM) and 1 mM of ATP. This mix was added on BgaA coated coverslips and binding events were imaged using a DIC microscope with 100X objective.
The instrument and techniques for optical trapping have been described in detail[41]. Protein conjugated beads were observed at room temperature in a custom-developed DIC microscope (Nikon TE2000-U) using a 100X, 1.4 numerical aperture oil objective. Images were acquired at video rate 30 frames/sec with a Cohu 4910 camera. Individual beads were trapped and brought in contact with the BgaA coated surface. The piezo stage was used to intermittently move the coverslip surface below the trapped bead at a constant velocity of 300 nm/s for 1 sec. The position of beads was tracked frame by frame using custom software.
Distance translocated by the p97 bead in the Z-axis was measured by first establishing a calibration curve with the bead being moved towards the slide surface at a 20 nm step size, which was estimated by plotting a greyscale histogram in ImageJ. The mean grey scale value was then estimated and normalized from initial bead position to final translocated bead position.

#### *In-vivo* sepsis model

All the experimental work on animals were done as per the guidelines of the Committee for the Purpose of Control and Supervision of Experiments on Animals (CPCSEA), India. The study protocol has been reviewed and approved by the Institutional Animal Ethics Committee (IAEC, Reg. no. 48/1999/CPCSEA) of Indian Institute of Science (IISc, Bengaluru, India) under the project number CAF/Ethics/380/2014. Briefly, 6-8 weeks old CD1 female mice with an average weight of 20 to 24 gm were injected intravenously via tail-vein injection with $10^6$ SPN. In two different cohorts, animals were treated intra-peritoneally with either DMSO (vehicle) or p97 inhibitor, NMS-873 (0.02 mg/kg). Mice were culled when they were moribund or at specific time points. Blood samples were obtained by cardiac puncture and excised spleens were processed via a handheld homogenizer. For enumeration of bacterial burden, blood and spleen homogenates were serially diluted and then plated on sheep blood agar plates, which were incubated overnight at 37°C, 5% $CO_2$, and bacterial colony numbers were assessed the following day.

### ELISA

Cytokine (IL-1β and IL-6) levels in the serum and spleen of mice infected with *SPN* or treated with lipopolysaccharide (LPS, 1 mg/kg) were measured using ELISA MAX™ Standard Set Mouse IL-6 (BioLegend 431301) and ELISA MAX™ Standard Set Mouse IL-1β (BioLegend 432601) according to the manufacturer's protocol.

### Immunohistochemistry

Spleens were fixed in 4% paraformaldehyde, embedded in paraffin and 5 μm thick paraffin sections were collected on poly-lysine coated glass slides. Tissue sections were incubated in 100% Xylene for 20 min, followed by washing in 1X PBS with 0.05% Tween 20. Staining was performed with anti-enolase Ab (specific for SPN, provided by Pro. Sven Hammarschmidt, Univ. of Greifswald, Germany; 1:5000) followed by AlexaFluor-488 conjugated secondary Ab (1:100). Sections were then washed with 1X PBS, mounted using Vecta-shield with DAPI (Vector Laboratories) and visualized under a Laser Scanning Confocal microscope (Carl Zeiss LSM 780).

### Statistical analysis

Statistical analyses were performed using GraphPad Prism software (version 9). Statistical tests undertaken for individual experiments are mentioned in the respective figure legends. Statistical significance was accepted at $p<0.05$. All multi-parameter analyses included corrections for multiple comparisons, and data are presented as means ± SD unless otherwise stated.

**Figure S1.**

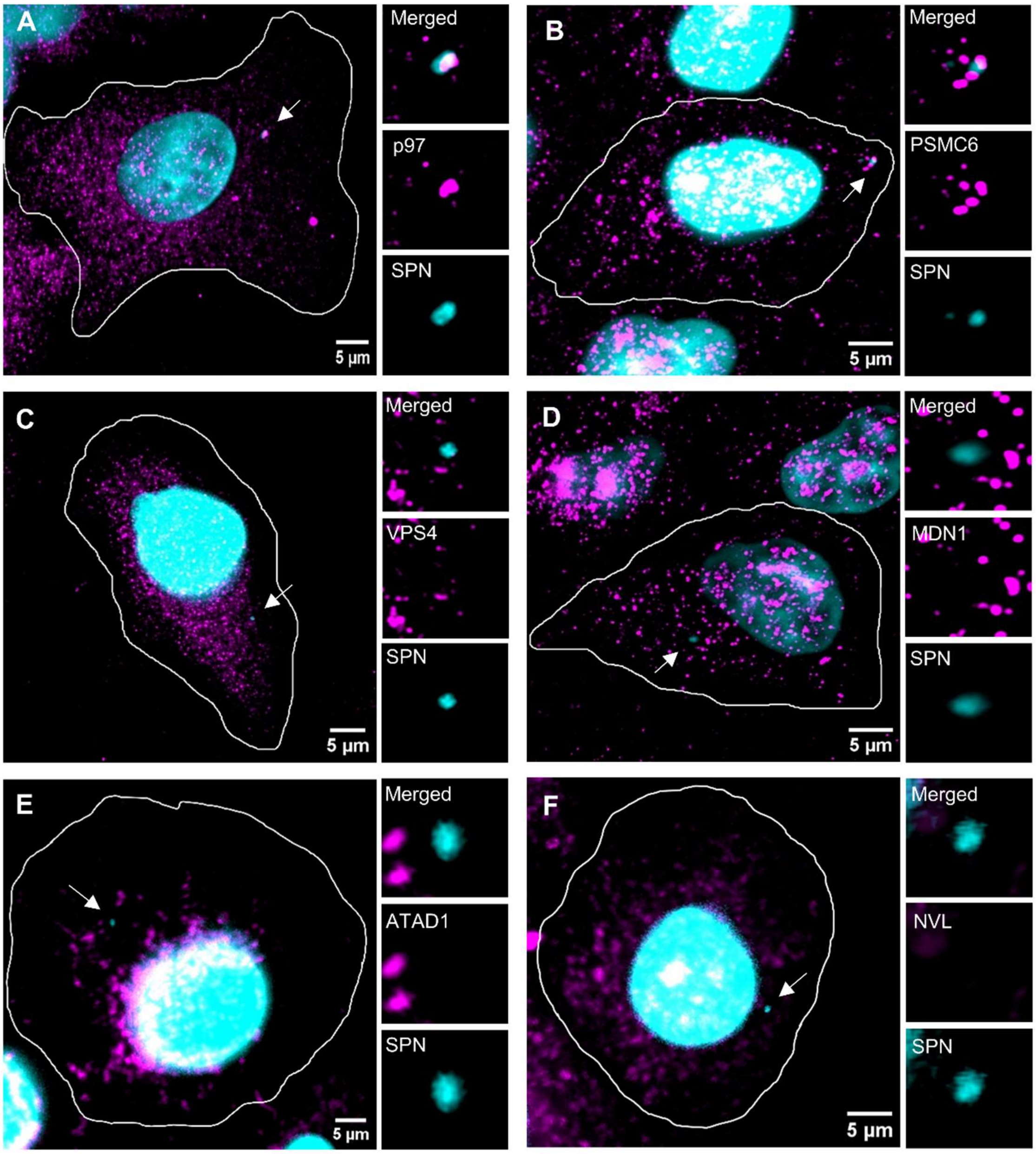


**Evaluation of association of different AAA-ATPases with intracellular SPN.** Structural Illumination Microscopy (SIM) showing intracellular SPN (cyan) in host cells with various AAA-ATPases (magenta). p97 **(A)**, PSMC6 **(B)**, VPS4 **(C)**, MDN1 **(D)**, ATAD1 **(E)** and NVL **(F)**. Arrows depict events shown in insets besides respective images. Scale bar, 5 µm. A549 cells were infected with SPN for 9 h. Pneumococcus was stained with DAPI and AAA-ATPases with respective primary Ab, followed by Alexa Fluor 488 conjugated to secondary Ab.

**Figure S2.**

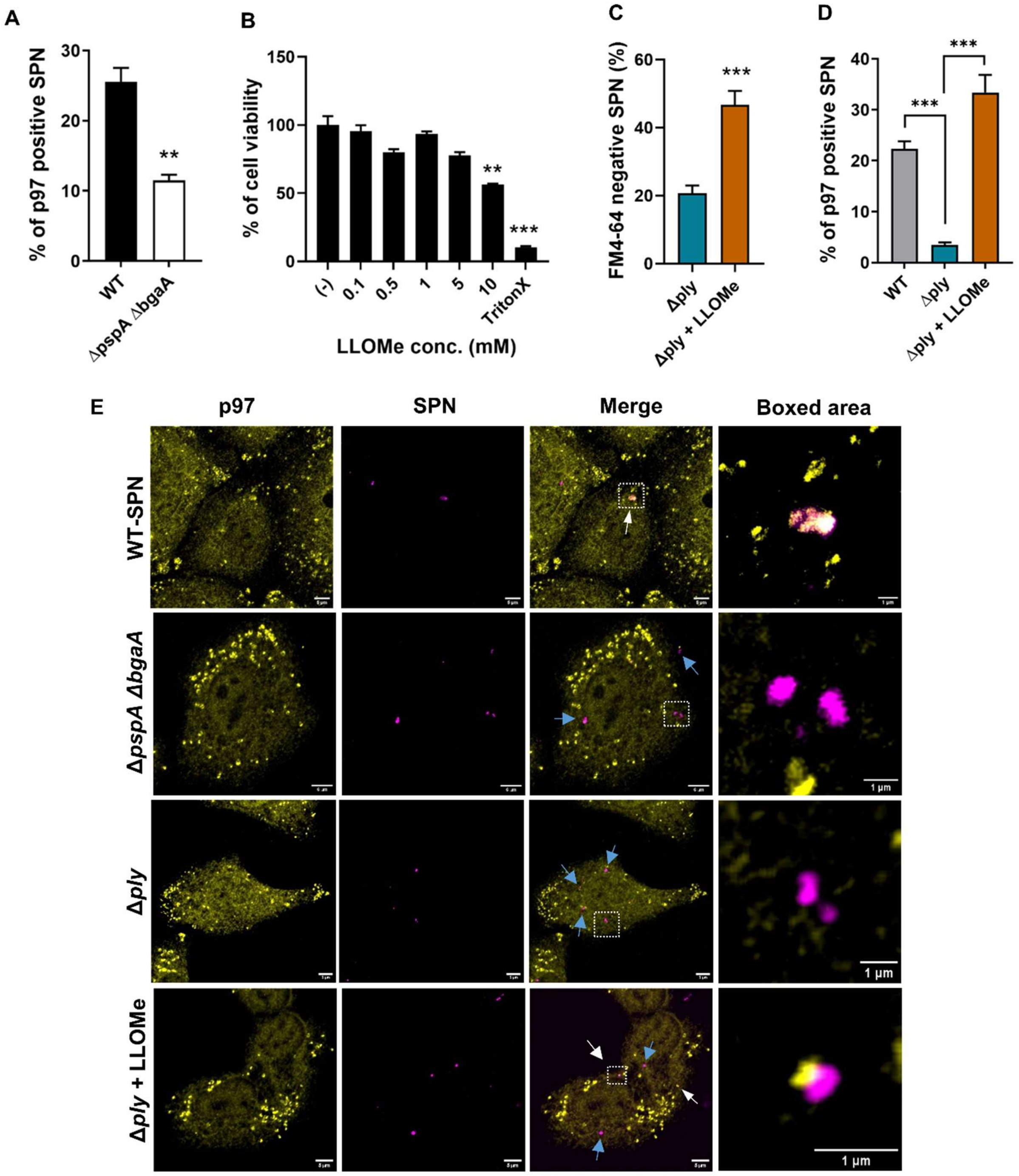


**Recruitment of p97 to the bacteria is endosome damage dependent. A.** Percentage of p97 association with WT SPN and BgaA and PspA knockout SPN (Δ*pspA* Δ*bgaA*) strains. n>100 events examined for three independent experiments. **B.** Percentage of viability of A549 cells upon treatment with different concentrations of endomembrane perforating agent L-leucyl-L-Leucine methyl ester (LLOMe, 3 mM). **C.** Percentage of FM4-64 (membrane staining dye) negative intracellular Δ*ply*

(Pneumolysin knock-out) SPN strain with or without treatment of host cells with LLOMe. n>100 events examined for three independent experiments. **D.** Percentage of p97's association with SPN lacking pneumolysin (Δ*ply*) and Δ*ply* mutant infected cells treated with LLOMe compared to WT SPN. n>100 events examined for three independent experiments. **E.** Confocal micrographs of host cells stained with p97 (Yellow) following infection with different variants of SPN (Magenta) (WT, Δ*pspA* Δ*bgaA*, Δ*ply*) with or without treatment with LLOMe. White arrows depict positive co-localization events, whereas sky-blue arrows designate SPN that did not associate with p97. Boxed areas in "Merged" panel are shown as enlarged images in the side. Scale bar, 5 µm. Statistical significance was assessed by one-way ANOVA followed by Dunnett's test in **(B-D).** Two-tailed unpaired student's t-test (nonparametric) was used for **(A).** $^{*}P < 0.05$; $^{**}P < 0.01$; $^{***}P < 0.005$. Data are means ± SD of N = 3 independent biological replicates.

**Figure S3.**

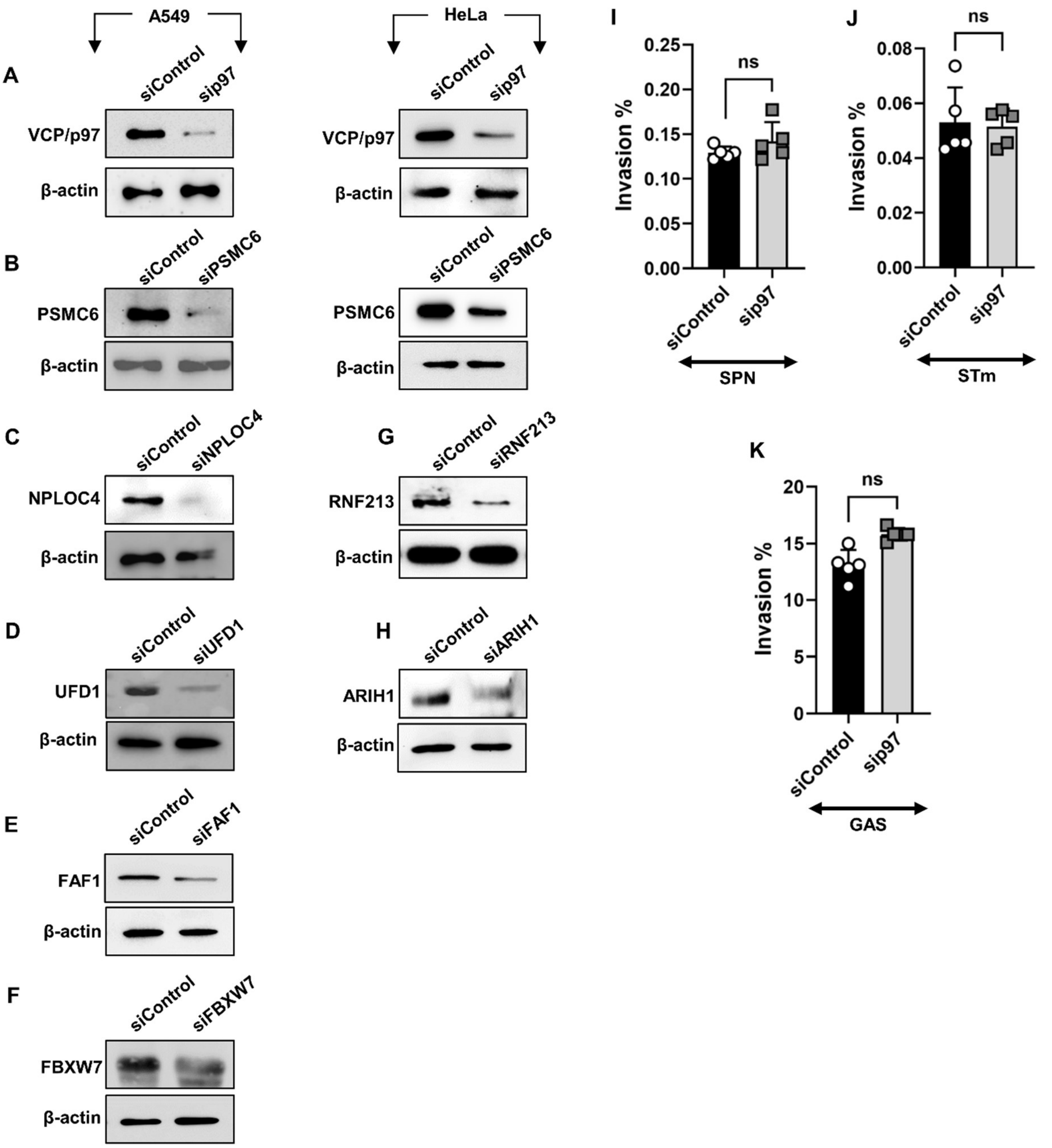


**siRNA mediated knockdown of gene expression and cellular invasion of SPN in p97 knockdown condition. A-B.** Immunoblot demonstrating the knockdown of p97 **(A)** and PSMC6 **(B)** in A549 and Hela cells. **C-F.** Immunoblot showing knockdown of NPLOC4 **(C),** UFD1 **(D)**, FBXW7 **(E)** and FAF1 **(F)** in A549 cells. **G-H.** Immunoblot showing knockdown of RNF213 **(G)** and ARIH1 **(H)** in Hela cells. **I-K.** Percentage of invasion of SPN **(I)**, STm **(J)** and GAS **(K)** in p97 downregulated cells compared to scrambled treated A549 cells. Mid-exponentially grown SPN, STm and GAS ($OD_{600nm}$

~ 0.4) were used to infect monolayers of A549s (with SPN at MOI ~ 10 and GAS at MOI ~ 2) or HeLa (with STm at MOI ~ 50) for 1 h. Following elimination of extracellular bacteria with antibiotics, bacterial invasion efficiency was calculated as: (recovered CFU/initial inoculum CFU) × 100%. Data are means ± SD of N = 4-5, independent biological replicates. Two-tailed unpaired student's t-test (nonparametric) was used for **(I-K)**. ns, non-significant.

**Figure S4.**

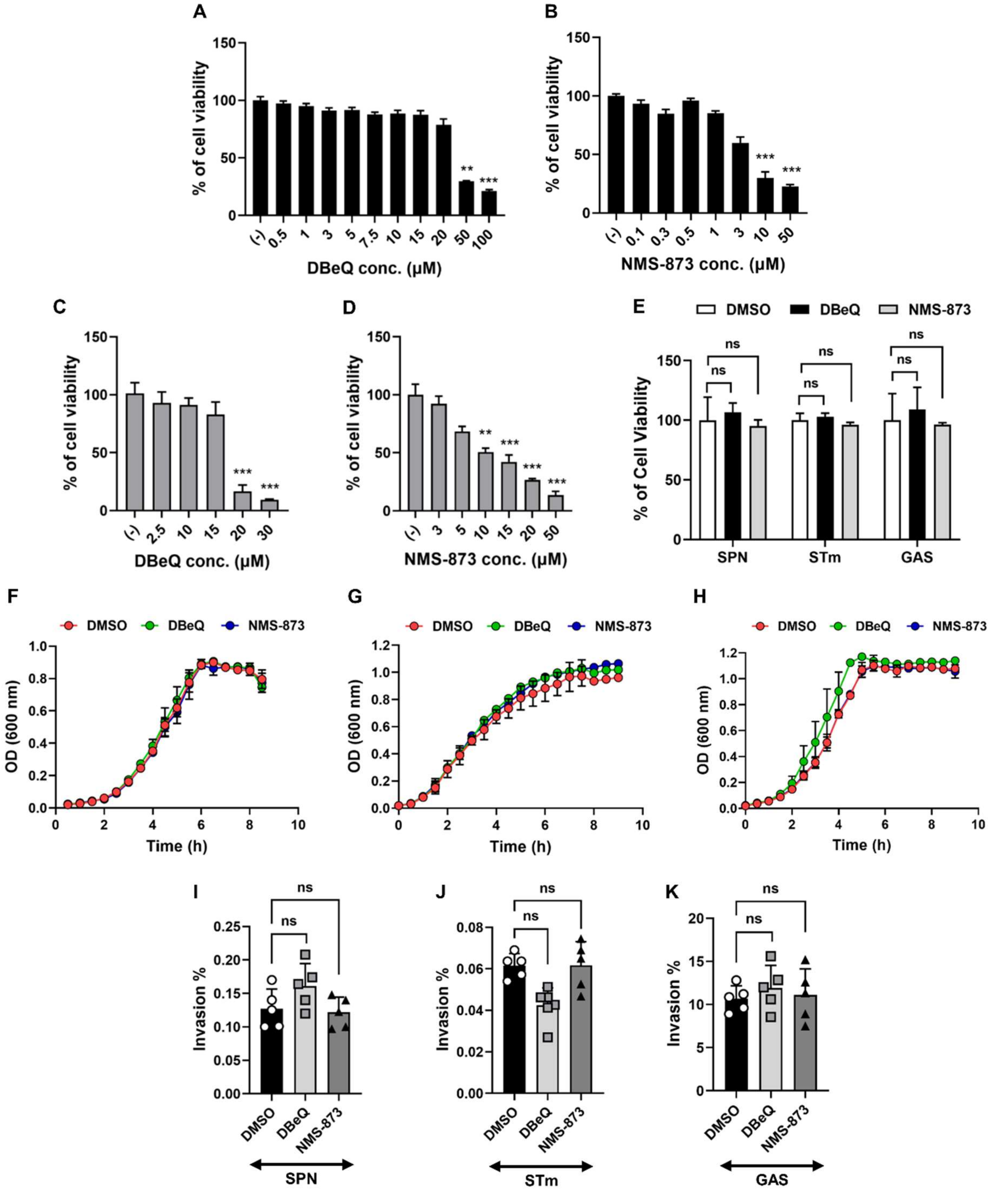


**Toxicity of p97 inhibitors and their effect on cellular invasion of pathogens. A-B.** Percentage of viability of A549 cells when treated with different concentrations of DBeQ **(A)** and NMS-873 **(B)**. **C-D.** Percentage of viability of HeLa cells when treated with different concentrations of DBeQ **(C)** and NMS-873 **(D)**. **E.** Percentage viability of host cells post-treatment with p97 inhibitors DBeQ (2.5 μM) and NMS-873 (1 μM) and subsequent infection with designated pathogens. A549 or HeLa cells were treated with vehicle (DMSO) or DBeQ or NMS-873 for 1 h prior to infection with SPN, STm or GAS. A549 cells were infected with SPN and GAS for 9 h while HeLa cells were infected with STm for 6 h. Post-infection, cells were assessed for viability by performing an MTT assay. Data are means ± SD of N = 3, independent biological replicates **(A-E)**. **F-H.** Graphs representing growth kinetics of SPN **(F)**, STm **(G)**, and GAS **(H)** when treated with DMSO or p97 inhibitors. The concentrations used, DBeQ (2.5 μM) and NMS-873 (1 μM), are similar to those used for cellular toxicity and invasion assays. Data are means ± SD of N = 3, independent biological replicates. **I-K.** Percentage of invasion of SPN **(I)** STm **(J)** and GAS **(K)** in host cells following treatment with p97 inhibitors, DBeQ (2.5 μM) and NMS-873 (1 μM). Bacterial invasion efficiency was calculated as: (recovered CFU/initial inoculum CFU) × 100%. Data are means ± SD of N = 5-6, independent biological replicates. Statistical significance was assessed by one-way ANOVA followed by Dunnett's test in **(A-E, I-K)**. ns, non-significant; **P < 0.01; ***P < 0.005.

**Figure S5.**

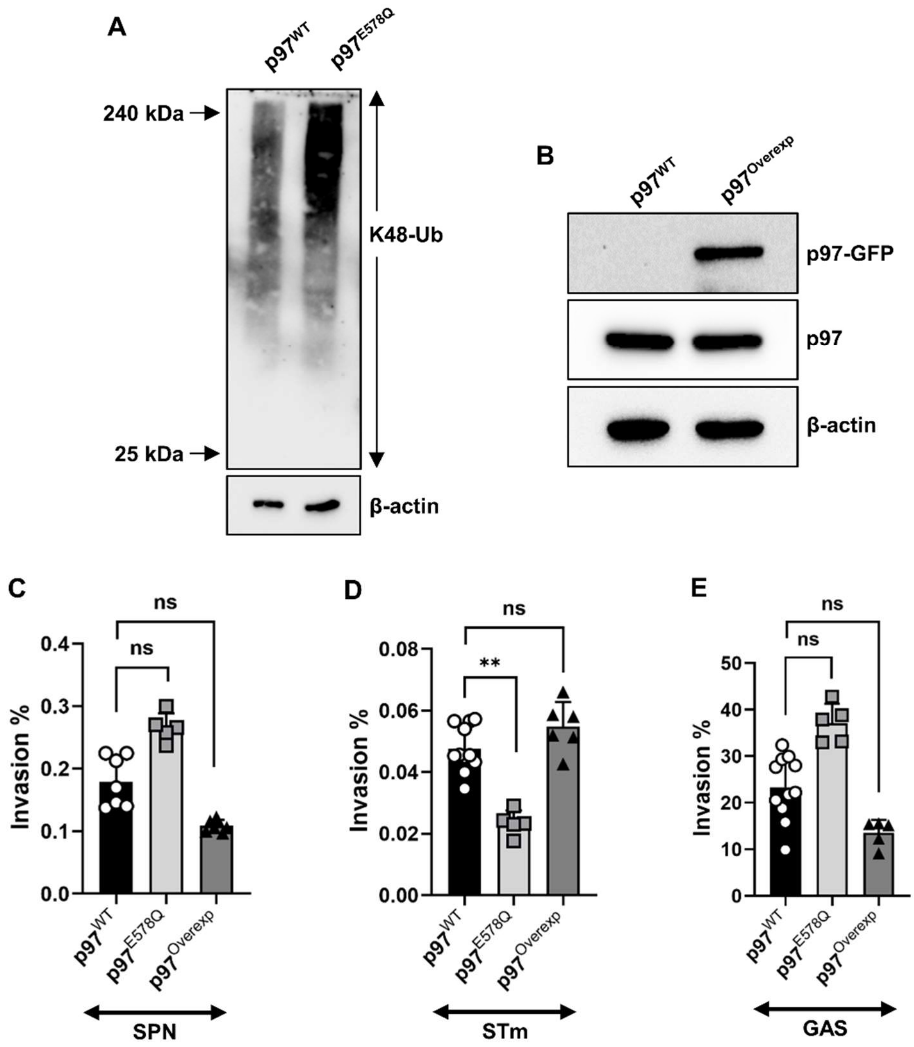


**Overexpression or expression of catalytically inactivated variant of p97 in host cells have unaltered effects on bacterial infections. A.** Immunoblot demonstrating the accumulation of K48-Ub substrates in the catalytically inactive p97$^{E578Q}$ cell line. Host cells were stably transfected with p97$^{E578Q}$ variant and presence of ubiquitinated proteins in cell lysates were detected by immunoblotting with anti-K48-Ub Ab. **B.** Immunoblot showing the expression of p97-GFP in host cells. Immunoblotting was performed on cell lysates with anti-GFP and anti-p97 Abs to detect expression of p97 as well as p97-GFP fusion proteins. β-actin was used as loading control. **C-E.** Percentage of invasion of SPN **(C)**, STm **(D)** and GAS **(E)** in host cells overexpressing p97 (p97$^{Overexp}$) or catalytically inactive form, p97$^{E578Q}$. Bacterial invasion efficiency was calculated as: (recovered CFU/initial inoculum CFU) × 100%. Data are means ± SD of N = 5-6, independent biological replicates. Statistical significance was assessed by one-way ANOVA followed by Dunnett's test in **(C-E)**. ns, non-significant; **P < 0.01.

**Figure S6.**

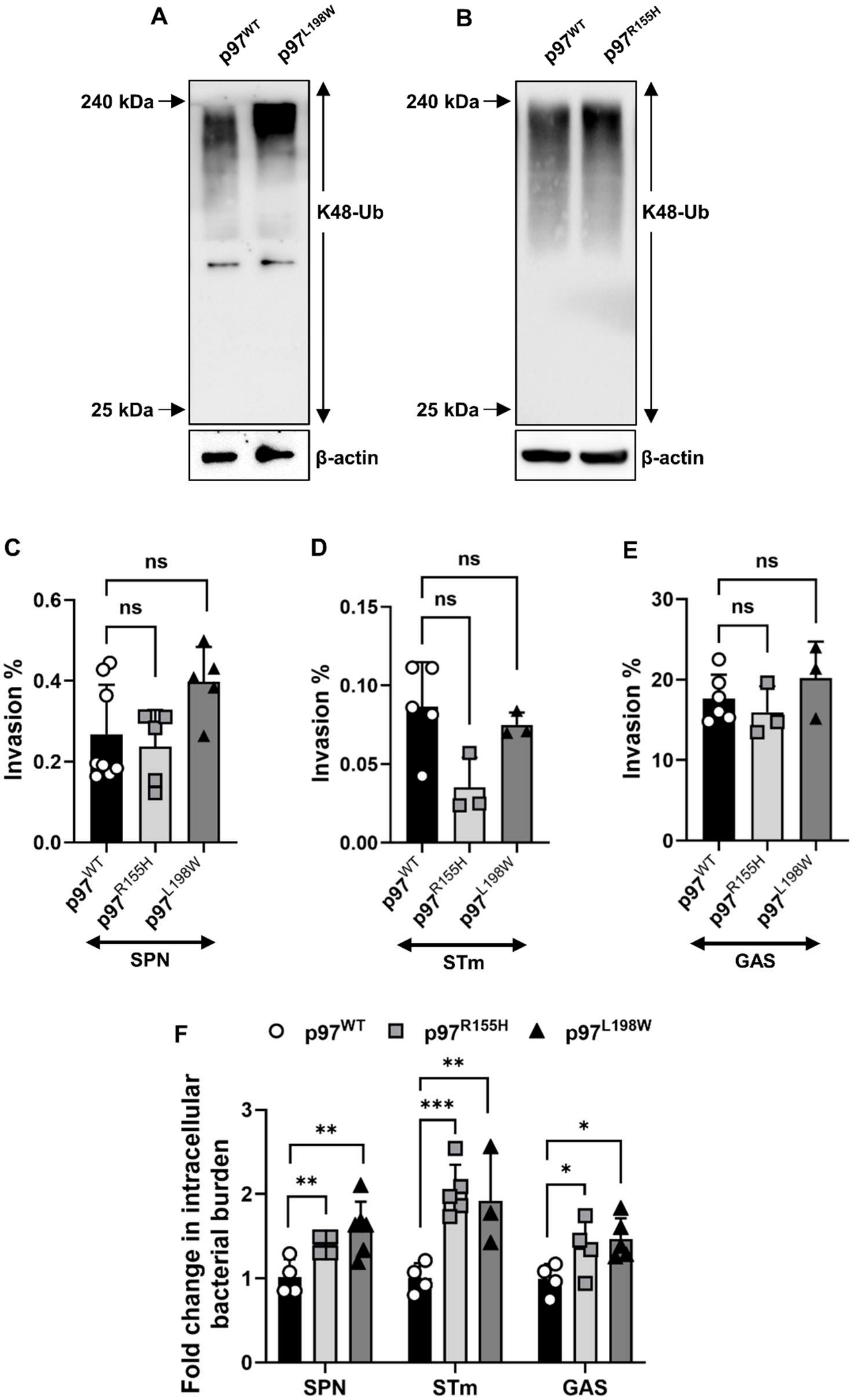
A
p97WT
p97L198W
240 kDa
25 kDa
K48-Ub
β-actin
B
p97WT
p97R155H
240 kDa
25 kDa
K48-Ub
β-actin
C
ns
ns
Invasion %
0.6
0.4
0.2
0.0
p97WT
p97R155H
p97L198W
SPN
D
ns
ns
Invasion %
0.15
0.10
0.05
0.00
p97WT
p97R155H
p97L198W
STm
E
ns
ns
Invasion %
30
20
10
0
p97WT
p97R155H
p97L198W
GAS
F
p97WT
p97R155H
p97L198W
Fold change in intracellular bacterial burden
3
2
1
0
**
**
**
***
*
*
SPN
STm
GAS

**Polymorphic mutations in p97 associated with multisystem proteinopathy render the host susceptible to bacterial infections. A-B.** Immunoblot demonstrating the accumulation of K48-Ub substrates in cells expressing polymorphic variants p97$^{L198W}$ **(A)** and p97$^{R155H}$ **(B)**. Host cells were stably transfected with different p97 variants and presence of ubiquitinated proteins in cell lysates was detected by immunoblotting with anti-K48-Ub Ab. **C-E.** Percentage of invasion of SPN **(C)**, STm **(D)** and GAS **(E)** in host cells expressing p97 polymorphic mutants, p97$^{L198W}$ and p97$^{R155H}$. Mid-exponentially grown SPN, STm and GAS ($OD_{600nm}$ ~ 0.4) were used to infect monolayers of A549s (with SPN at MOI ~ 10 and GAS at MOI ~ 2) or HeLa (with STm at MOI ~ 50) stably transfected with p97$^{L198W}$ or p97$^{R155H}$ variants for 1 h. Following elimination of extracellular bacteria with antibiotics, bacterial invasion efficiency was calculated as: (recovered CFU/initial inoculum CFU) × 100%. Data are means ± SD of N = 5-6, independent biological replicates. **F.** Graph representing fold change in intracellular burden of different bacteria in p97 mutant host cells compared to p97$^{WT}$. A549s are infected with SPN (MOI ~ 10) and GAS (MOI ~ 2) for 9 h while HeLa cells were infected with STm (MOI ~ 50) for 6 h. At these time points intracellular bacterial numbers were deciphered and fold change in intracellular bacterial burden is calculated as ratio of intracellular bacterial CFU of the test (mutant cell lines) to that of control (p97$^{WT}$ cells). Data are means ± SD of N = 4-5, independent biological replicates. Statistical significance was assessed by one-way ANOVA followed by Dunnett's test in **(C-F)**. ns, non-significant; *P < 0.05; **P < 0.01; ***P < 0.005.

**Figure S7.**

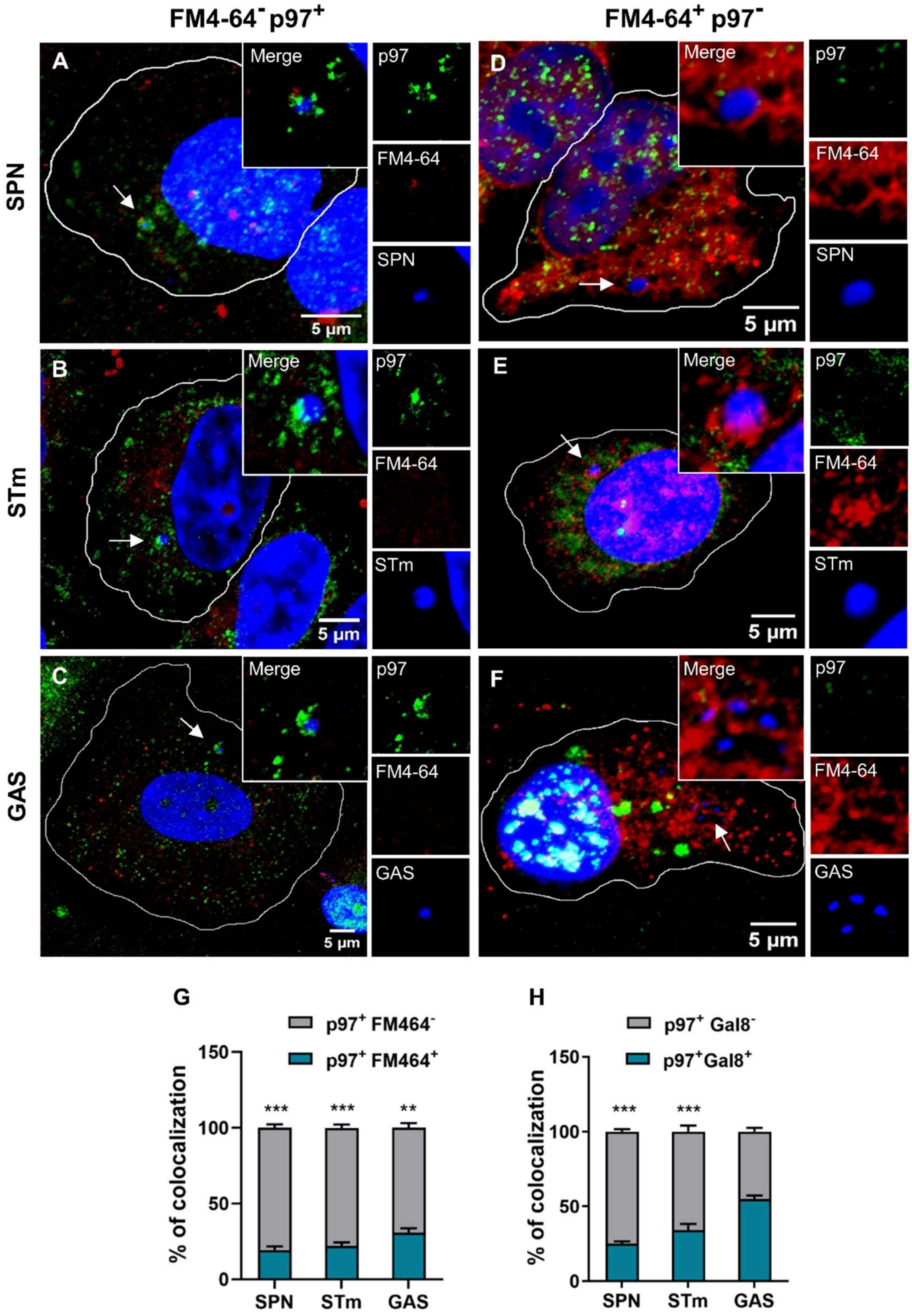


**Cytosolic SPN is targeted by p97. A-C.** Confocal micrographs showing p97 (Green) positive SPN **(A)**, STm **(B)**, and GAS **(C)** (Blue) are devoid of membrane marker FM4-64 (Red). **D-F.** Confocal images depicting FM4-64 (Red) positive SPN **(D)**, STm **(E)**

and GAS **(F)** that fails to recruit p97 (Green). Arrows depict events shown in insets besides respective images.  Scale bar, 5 µm. **G-H.** Percentage population of p97-positive bacteria that are marked with or without membrane marker FM4-64 **(G)** or damaged endosome marker Gal8 **(H)**. n>50 events examined for three independent experiments. A549 and Hela cells were infected with SPN, GAS for 9 h and STm for 6 h, respectively. Bacteria were stained with DAPI, membranes were marked with FM4-64, while p97 and Gal8 were with respective primary Ab, followed by Alexa Fluor 488 or Alexa Fluor 555 conjugated to secondary Ab. Statistical significance was assessed by one-way ANOVA followed by Dunnett's test in **(G, H)**. ***P < 0.005.

**Figure S8.**

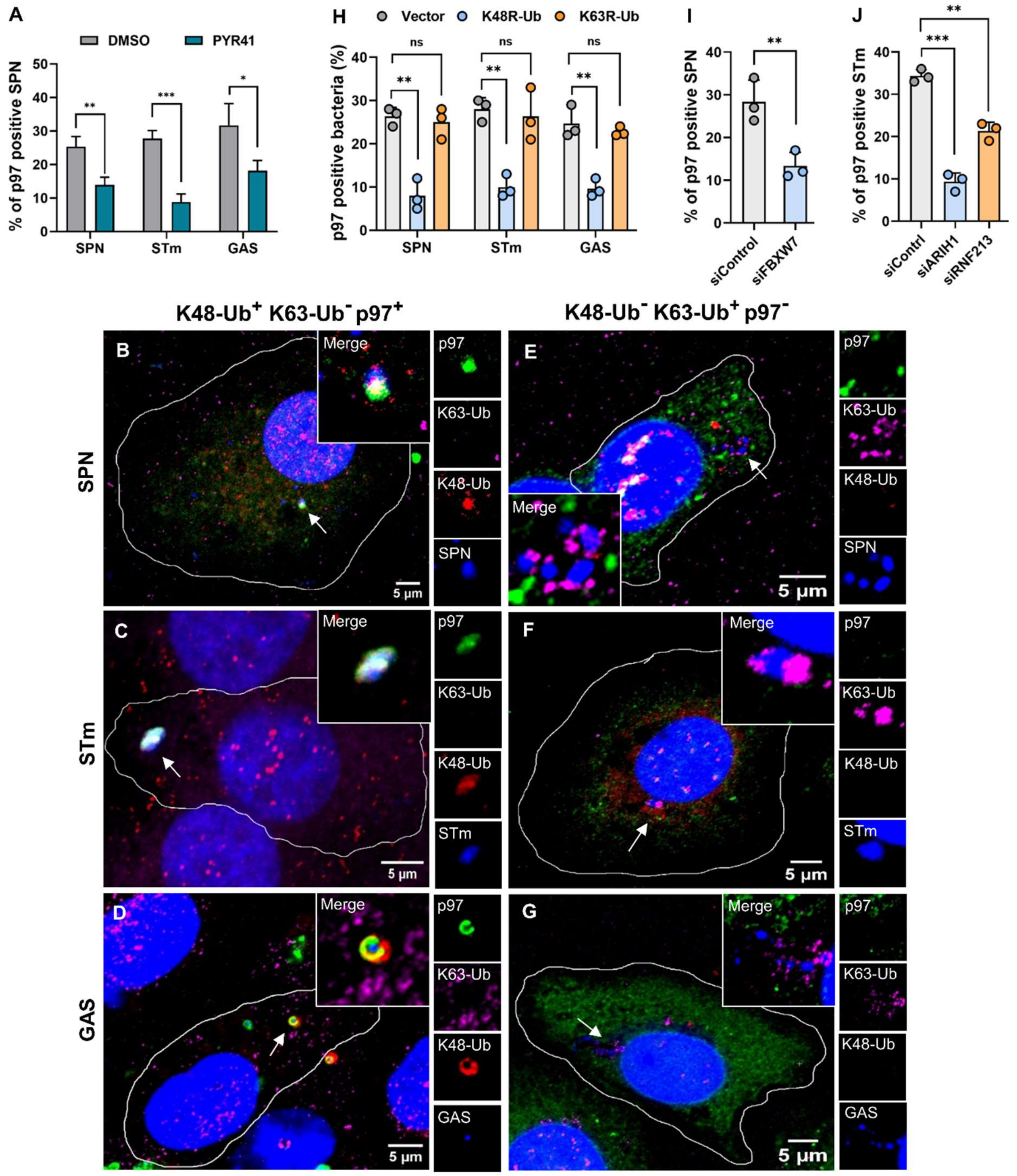


**K48-polyubiquitinated bacteria recruits p97. A.** Percentage of p97-positive bacteria in host cells treated with E1-Ub ligase inhibitor PYR41 (10 µM) compared to DMSO control. Cells were treated with PYR41 for 1 h before being infected with different pathogens. At 6-9 h, post-infection, association of intracellular bacteria (stained with DAPI) with p97 (stained with anti-p97 Ab) was assessed by quantifying confocal images. n>100 events examined for three independent experiments. **B-D.** Confocal micrographs showing the association of p97 (Green) positive SPN **(B)**, STm **(C)**, and GAS **(D)** (Blue) with K48-Ub (Red) but not with K63-Ub (Magenta). **E-G.** Confocal images depicting K63-Ub (Magenta) positive but K48-Ub (Red) negative SPN **(E)**, STm **(F)** and GAS **(G)** that fail to recruit p97 (Green). Arrows depict events shown in insets besides respective images. Bacteria were stained with DAPI, p97 was conjugated to GFP, K48-Ub and K63-Ub were stained with respective primary Ab, followed by Alexa Fluor 555 or Alexa Fluor 633 conjugated to secondary Ab. Scale bar, 5 μm. **H.** Percentage of p97-positive bacteria in cells expressing mutant ubiquitin (K48R and K63R) compared to wild-type cells. Defects in ability of mutant ubiquitin variant expressing host cells to conjugate specific ubiquitin chain topologies, without hampering conjugation of other ubiquitin chain types to cellular substrates were assessed earlier[1]. n>100 events examined for three independent experiments. **I.** Percentage of association of p97 with SPN in FBXW7 downregulated host cells. FBXW7 expression in A549s were knocked down by siFBXW7, followed by infection with SPN. n>100 events examined for three independent experiments. **J.** Percentage association of STm with p97 in siARIH1 and siRNF213 treated HeLa cells. While AIRH1 is reported to ubiquitinate cytosolic STm with K48-Ub chain types, RNF213 ubiquitinates LPS present on STm outer membrane[2, 3]. n>100 events examined for three independent experiments. Statistical significance was assessed by one-way ANOVA followed by Dunnett's test in **(A, H)** or two-tailed unpaired student's *t*-test (nonparametric) **(I, J)**. ns, non-significant; *P < 0.05; **P < 0.01; ***P < 0.005.

**Figure S9.**

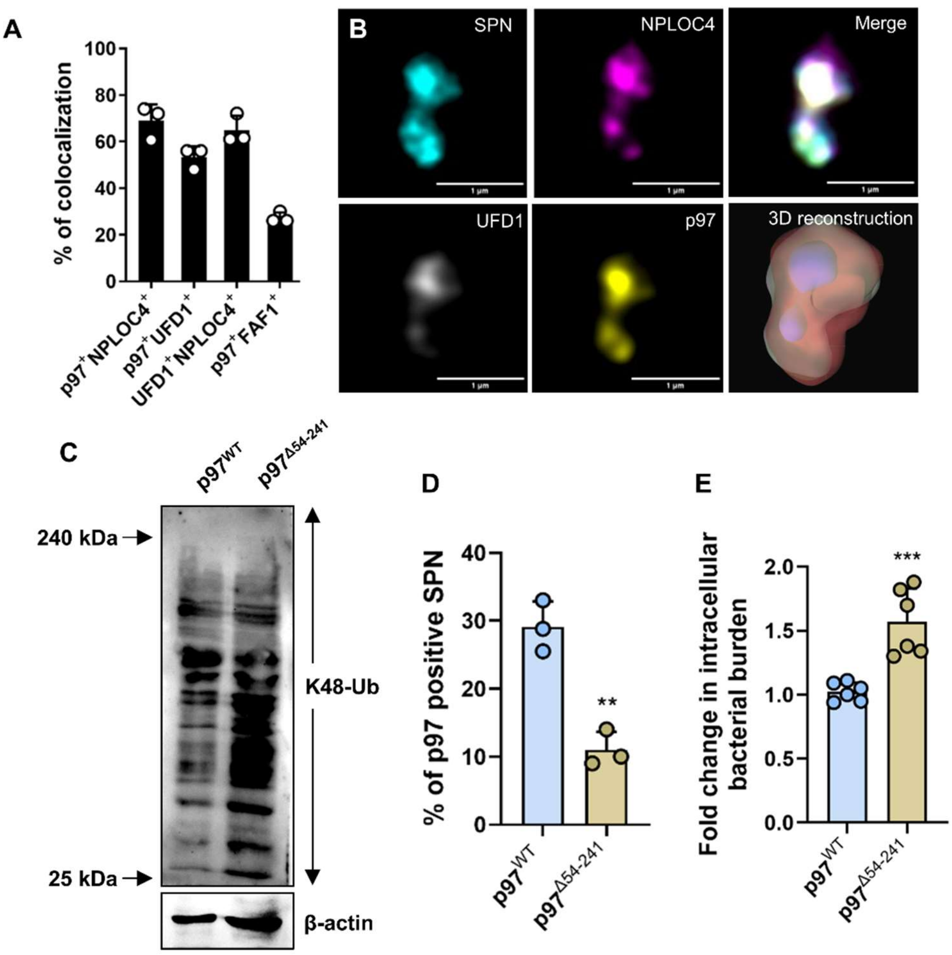


**NPLOC4 and UFD1 are the primary cofactors involved in p97's targeting of bacteria. A.** Percentage of p97 positive intracellular SPN associating with various cofactors, such as, NPLOC4, UFD1, FAF1 or NPLOC4 positive SPN associating with UFD1. n>50 events examined for three independent experiments. **B.** Structural illumination microscopy (SIM) showing the co-localization of SPN (Cyan) with UFD1 (White), NPLOC4 (Magenta) and p97 (Yellow). Scale bar, 1 µm. 3D reconstruction of the merged image using IMARIS is also represented. Bacteria were stained with DAPI, p97 was conjugated to GFP, NPLOC4 and UFD1 were stained with respective primary Ab, followed by Alexa Fluor 633 or 555 conjugated to secondary Ab. **C.** Immunoblot showcasing the accumulation of K48-Ub substrates in host cells expressing p97$^{\Delta 54\text{-}241}$. Host cells were stably transfected with truncated p97 variant and inability to clear ubiquitinated proteins was detected by immunoblotting with anti-K48-Ub Ab. **D.** Quantitative analysis showcasing association of p97 with intracellular SPN in host cells expressing truncated p97$^{\Delta 54\text{-}241}$ compared to wild-type. n>100 bacteria/coverslip. **G.** Fold change in intracellular burden of SPN in host cells expressing truncated p97$^{\Delta 54\text{-}241}$ compared to wild-type cells. Host cells are infected with SPN (MOI ~ 10) for 9 h and fold change in intracellular bacterial burden is depicted as ratio of intracellular bacterial CFU of the test to that of control at specified time point. Data are means ± SD of N = 6, independent biological replicates. Statistical significance was assessed by Two-tailed unpaired student's *t*-test (nonparametric) **(D, E)**. **P < 0.01; ***P < 0.005.

**Figure S10.**

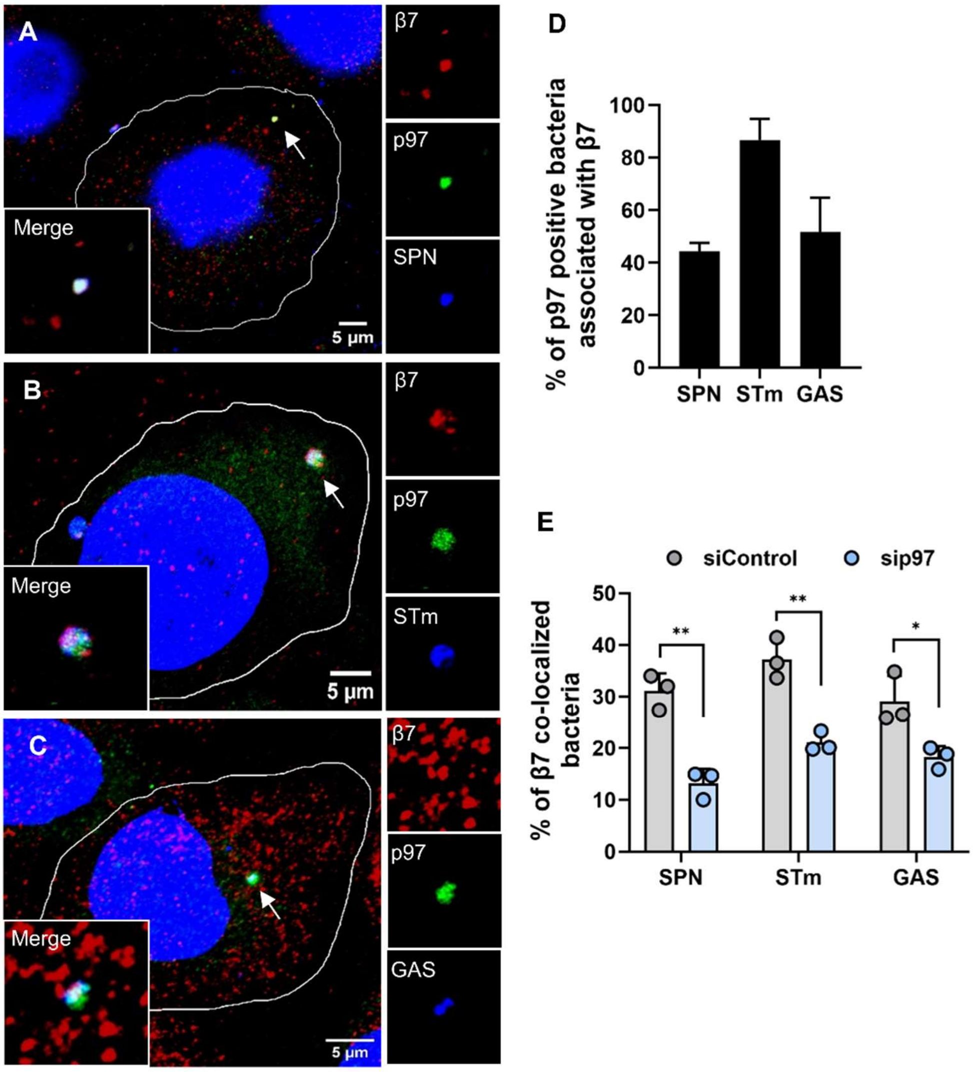


**Coordination of p97 with proteasome is required for bacterial targeting. A-C.** Confocal micrographs showing the association of p97 (Green) positive SPN **(A)**, STm **(B)**, and GAS **(C)** (Blue) with proteasomal marker, β7 (Red). Arrows depict events shown in insets besides respective images. Scale bar, 5 µm. **D.** Percentage association of p97-positive pathogens with the proteasome (marked by β7). n>100 events examined for three independent experiments. **E.** Percentage of β7 (Proteasomal marker) positive intracellular bacteria in host cells treated with sip97 compared with siControl (Scramblase). A549 and Hela cells were infected with SPN, GAS for 9 h and STm for 6 h, respectively. Bacteria were stained with DAPI, p97 and β7 with respective primary Ab, followed by Alexa Fluor 488 or Alexa Fluor 555 conjugated to secondary Ab. n>100 events examined for three independent experiments. Statistical significance was assessed by one-way ANOVA followed by Dunnett's test in **(E)**. *P < 0.05; **P < 0.01.

**Figure S11.**

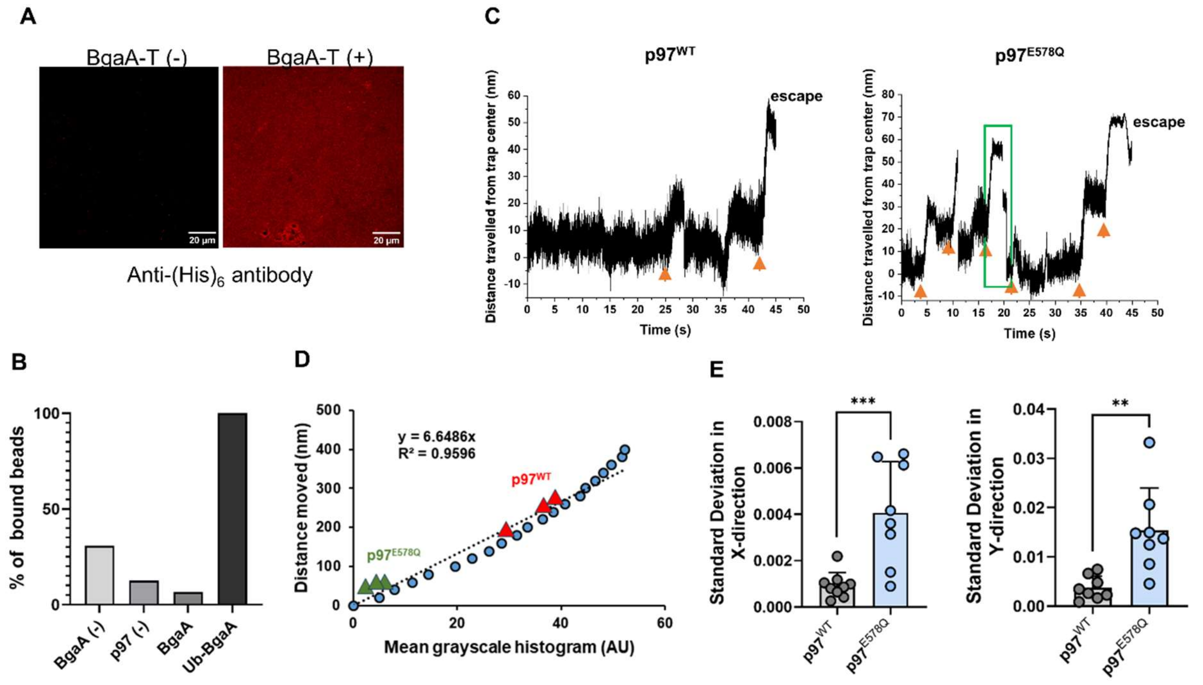


**p97 unfolds and translocates along BgaA-T. A.** Immunostaining of BgaA-T (a truncated version of BgaA, consisting of amino acids 1 to 1049) deposited on a hydrophilic coverslip using anti-(His)$_6$ antibody. Scale bar, 20 μm. **B.** Beads (conjugated or unconjugated with protein) were trapped and brought close to the coverslip surface. The percentage of beads that bound the surface was calculated and plotted for each of the conditions described in the figure. n>20 beads. **C.** Distance from trap center versus time plot the trap center giving multiple force generating events (one of which is depicted in the green box). Representative beads conjugated with p97$^{WT}$ (Left) and p97$^{E578Q}$ (Right) are shown. Orange arrows indicate the time points at which piezo was moved. On moving the piezo, the p97$^{E578Q}$ mutant binds the surface but keeps falling back to such events are few. Once the piezo is moved, the bead binds the surface and extracts the protein hence escape the trap (as shown). **D.** Streptavidin bead was moved in 20 nm steps in + and - Z positions and images were acquired at each step. A calibration curve – position versus intensity of greyscale histogram, was generated using the obtained values to calculate displacement for protein-conjugated beads, when they bind to BgaA surface. Blue dots represent intensity values obtained with only bead. Red triangles and green triangles are values obtained with p97 and $p97^{E578Q}$ beads, respectively. **E.** Graph showing the standard deviation in the X direction (left) and Y direction (right) for both p97 and $p97^{E578Q}$ beads bound to BgaA surface. Two-tailed unpaired student's t-test (nonparametric) was used for **(E)**. **P < 0.01; ***P < 0.005.

**Figure S12.**

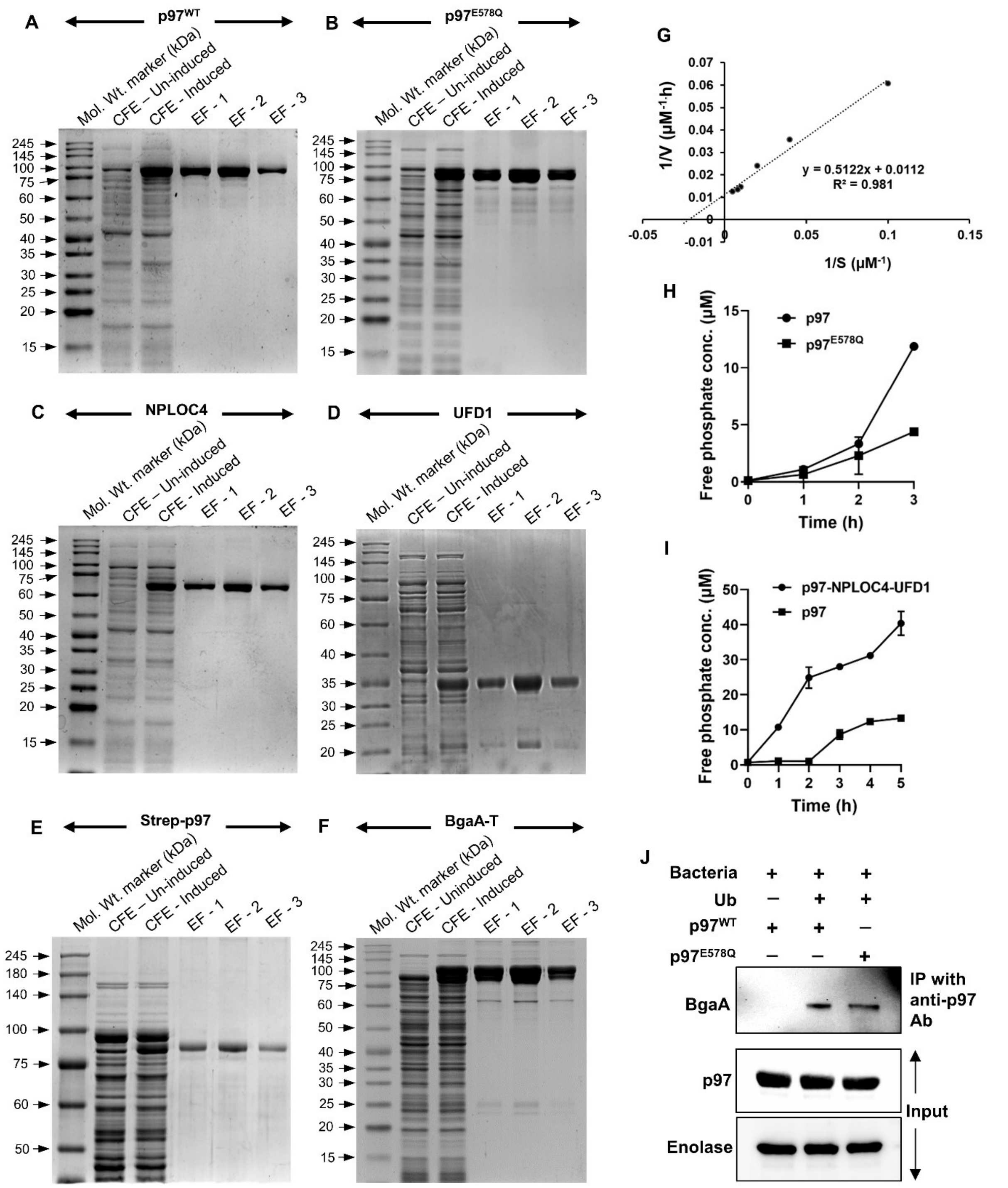


**Purification of proteins for *in-vitro* ubiquitination and ATPase assay. A-F.** SDS PAGE of p97 **(A)**, p97$^{E578Q}$ **(B)**, NPLOC4 **(C)**, UFD1 **(D)**, Strep-p97 **(E)**, BgaA-T **(F)** purified using Ni-NTA column from cell lysate of *E. coli* cells expressing the proteins. Molecular weight of markers is mentioned in the side of the SDS gels. CFE: Cell-free extract; EF: Elution fraction. *E. coli* Bl21(DE3) strain was used for expression of p97 or p97 variants and BgaA-T, while *E. coli* Rosetta(DE3) was used for expression of NPLOC4 and UFD1. Protein expression was induced using IPTG (0.2 mM for p97 or its variants and 1 mM NPLOC4 and UFD1). **G.** Lineweaver-Burk plot of purified p97 (1 μM) as measured following estimation of released phosphate from ATP (50 μM) by Malachite Green Phosphate assay. **H-I.** Comparison of the ATPase activity between $p97^{WT}$ and catalytic mutant $p97^{E578Q}$ **(H)** or p97 with or without its cofactors, UFD1 and NPLOC4 **(I). J.** Immunoblot showcasing the interaction of purified $p97^{WT}$ and $p97^{E578Q}$ to SPN surface protein BgaA upon incubating 5 μM p97 with $10^7$ SPN cells, followed by immunoprecipitation with anti-p97 antibody. Presence of BgaA in the immune-precipitated fraction was determined by immunoblotting with anti-BgaA Ab.

**Figure S13.**

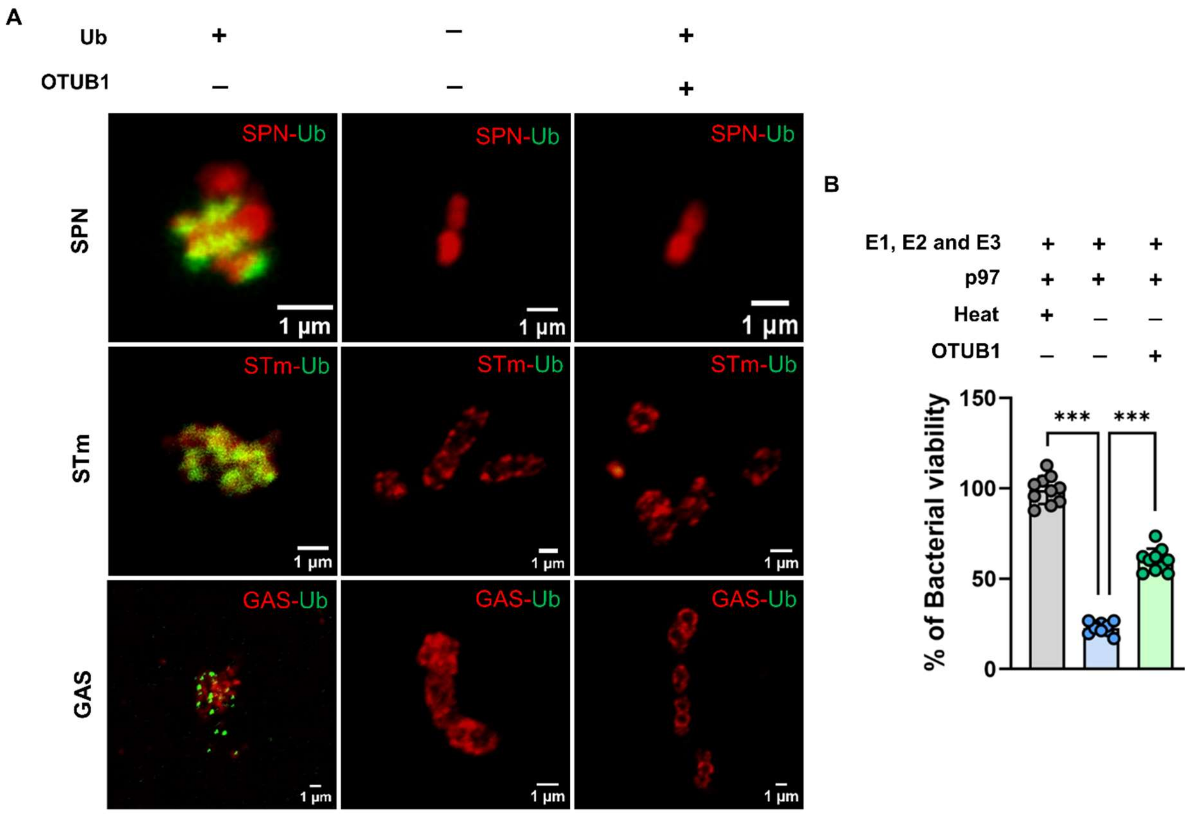


***In-vitro* ubiquitination of bacteria. A.** Confocal microscopy images demonstrating ubiquitination of bacteria in an *in-vitro* reconstituted reaction. $10^7$ bacterial cells/ml were incubated with cell lysate (2 mg/ml total protein) and ATP (1 mM) for 1 h at room temperature and ubiquitination of bacteria was analyzed by immunofluorescence following treatment with anti-K48-Ub Ab. Ubiquitin and bacterial surface-specific markers were stained with respective primary Ab, followed by Alexa Fluor 488 or Alexa Fluor 555 conjugated to secondary Ab. Scale bar, 1 µm. **B.** Percentage of bacterial viability upon treatment of ubiquitinated (by UBE1, UbE2C, and Rbx1-Skp1-Cul1-Fbxw7) or deubiquitinated bacteria (OTUB1 treated) with active p97 complex compared to denatured complex. Data are means ± SD of N = 10, independent biological replicates. Statistical significance was assessed by one-way ANOVA followed by Dunnett's test in **(B)**. ***P < 0.005.

**Figure S14.**

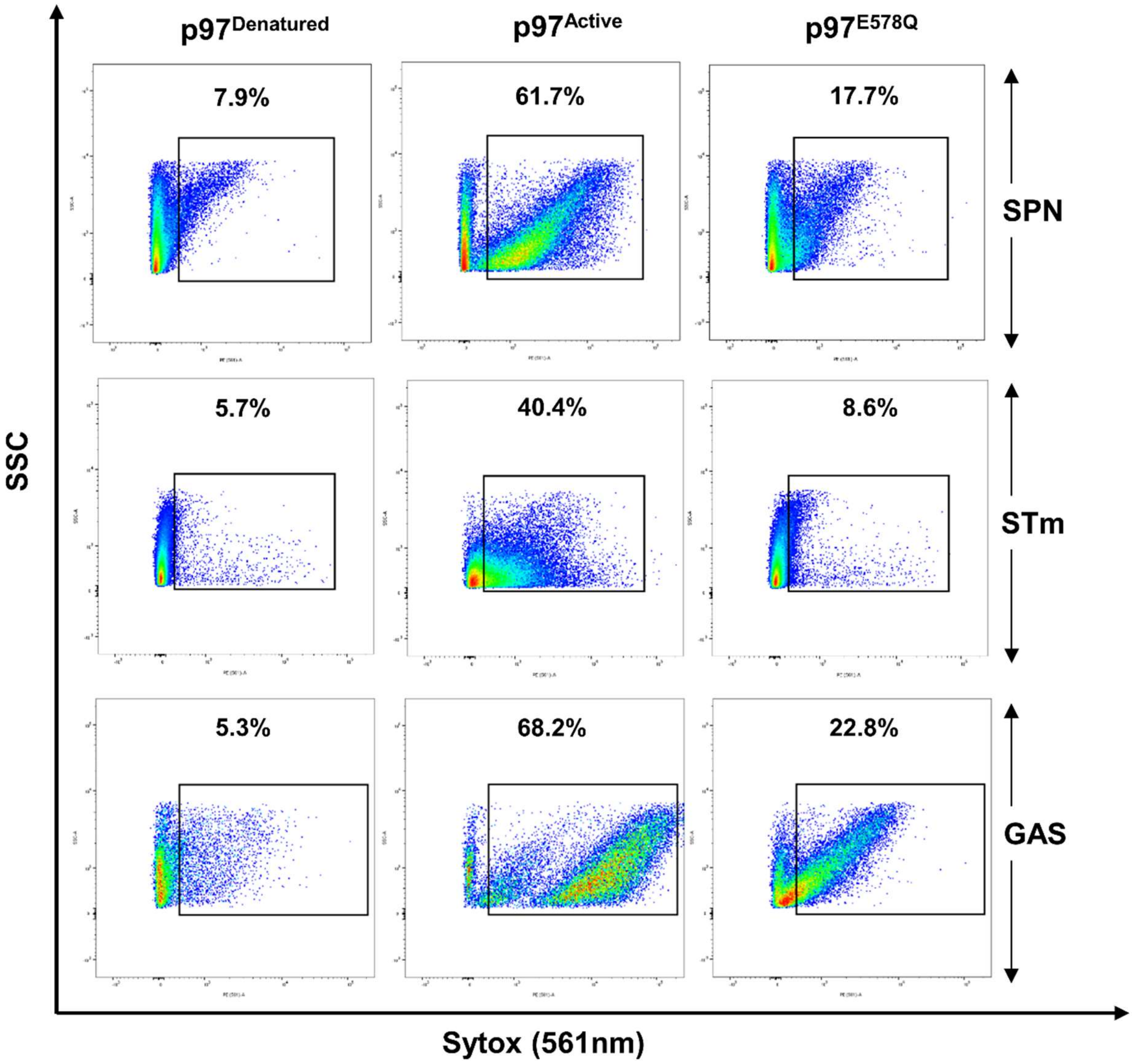


**p97 compromises bacterial membrane.** FACS analysis of Sytox internalization in SPN, STm and GAS upon treatment with active p97 complex compared to denatured p97 control ($p97^{Denatured}$) and $p97^{E578Q}$. >40000 events were analysed. Data was plotted using FlowJo® software.

**Figure S15.**

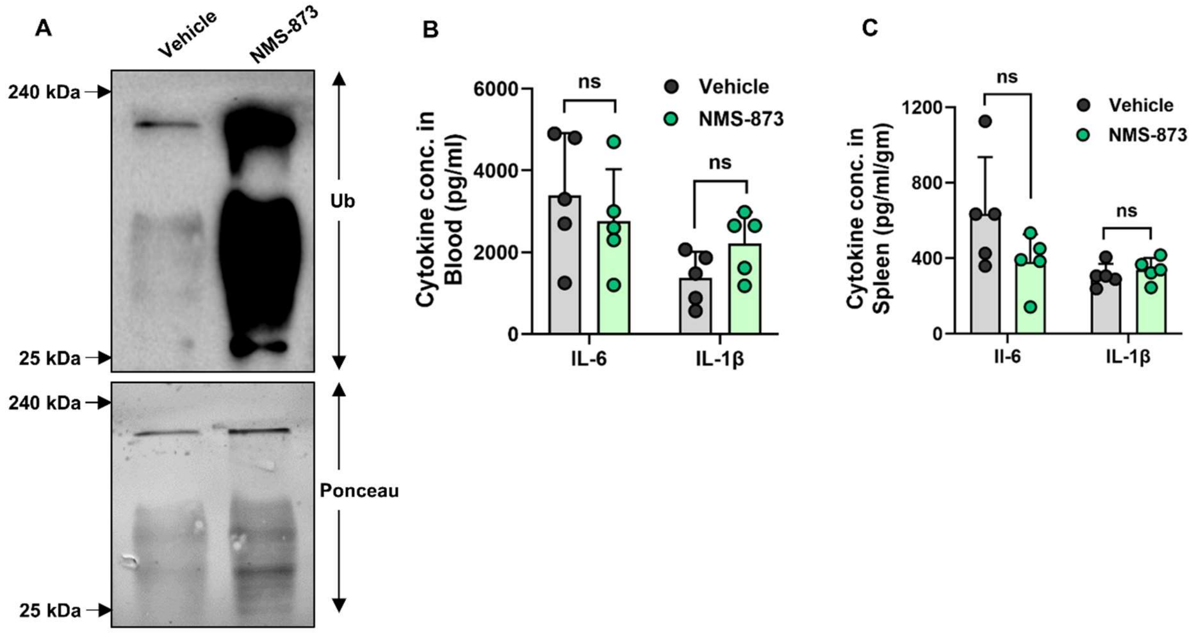


**NMS-873 inhibits p97 and does not hamper mice immunity. A.** Immunoblot demonstrating the accumulation of K48-Ub in spleen tissues of mice upon treatment with NMS-873 (0.02 mg/kg). **B-C.** Cytokine release in blood **(B)** and spleen **(C)** in mice at 12 h post administration of LPS (1 mg/kg). 1 h prior to LPS administration animals were treated with vehicle (DMSO) or NMS-873. n=5 animals per group. Statistical significance was assessed by two-way ANOVA followed by Bonferroni test. ns, non-significant.

**Table S1: List of strains.**

| Strain | Source |
|---|---|
| *Streptococcus pneumoniae* R6 (Serotype 2) | Prof. TJ Mitchell, Univ. of Birmingham, UK |
| *Streptococcus pneumoniae* D39 (Serotype 2) | Prof. TJ Mitchell, Univ. of Birmingham, UK |
| *Salmonella enterica subsp. enterica serovar* Typhimurium ATCC 14028 | ATCC |
| *Streptococcus pyogenes* Strain JRS4 | Prof. Victor Nizet, University of California, San Diego School of Medicine |
| SPN Δ*ply* | Surve *et al.,* 2018 |
| SPN Δ*bgaA* Δ*pspA* | Apte *et al.,* 2023 |

**Table S2: List of plasmids.**

| Plasmid Name | Details |
|---|---|
| pAB709 | pMRX-EGFP-p97$^{E578Q}$, Amp$^{R}$ |
| pAB712 | pMRX-EGFP-p97$^{WT}$, Amp$^{R}$ |
| pAB713 | pET28a-p97, Kan$^{R}$ |
| pAB720 | pET41b-NPLOC4-(His)$_{6}$, Kan$^{R}$ |
| pAB721 | pMRX-p97$^{R155H}$, Amp$^{R}$ |
| pAB722 | pMRX-p97$^{L198W}$, Amp$^{R}$ |
| pAB730 | pET28a- p97$^{E578Q}$, Kan$^{R}$ |
| pAB733 | pMRX-p97$^{54-241}$, Amp$^{R}$ |
| Addgene#117107 | pET41b-UFD1, Kan$^{R}$ |
| pAB1003 | pET28a-p97-FtsHp, Kan$^{R}$ |

**Table S3: List of Primers.**

| Name | Primer Sequence (5' to 3') |
|---|---|
| pMRX-p97$^{E578Q}$-F | ATCTGCCACCATGGTGAGCAAGGGCGAG |
| pMRX-p97$^{E578Q}$-R | CGAGGTGCGGCCGCTTATCTAGATCCGGTGGA |
| pMRX-p97$^{WT}$-F | CTTCTTTGATGAGTTAGATTCAATTGCCAAGGCT |
| pMRX-p97$^{WT}$-R | ATCTAACTCATCAAAGAAGAGTACACAGGGGG |
| pET28a-p97-F | ATTACACCATGGCCTCTGGAGCCGATTC 3 |
| pET28a-p97-R | CTAAATCTCGAGTCTAGATCCGGTGGATCCGC |
| pET41b-NPLOC4-F | ATTAATCCATGGTGGCCGAGAGCATCATAATT |
| pET41b-NPLOC4-R | ATTAATGCGGCCGCGGTCCTGGGGAGGCTGCACATCTC GCA |
| pMRX-p97$^{R155H}$-F | TTTCCTTGTCCACGGTGGGATGCGTGCTGT |
| pMRX-p97$^{R155H}$-R | CCACCGTGGACAAGGAAAATATCTCCTTTA |
| pMRX-p97$^{L198W}$-F | GAGTCCTGGAATGAAGTAGGCTATGATGAC |
| pMRX-p97$^{L198W}$-R | TACTTCATTCCAGGACTCCTCCTCATCCTCTC |
| pET28a-p97$^{E578Q}$-F | ATTACACCATGGCCTCTGGAGCCGATTC |
| pET28a-p97$^{E578Q}$-R | CTAAATCTCGAGTCTAGATCCGGTGGATCCGC |
| pMRX-p97$^{54\text{-}241}$-F | CTATATGGACCTCCTGGGACA |
| pMRX-p97$^{54\text{-}241}$-R | ACCTCTGAACAACTGTAGTTC |
| pET28a-p97-FtsHp-F | GCATTAGGATCCACCGGATCTAGAATGGTGATGACGGA AGCGCAGAAAG |
| pET28a-p97-FtsHp-R | GAAGTACTCGAGCTTGTCGCCTAACTGCTCTGACATGG |

**Movie S1. Molecular dynamic simulation of BgaA pulling by p97.** Steered molecular dynamic (SMD) simulation of p97 mediated pulling of BgaA, covalently attached to peptidoglycan surface. Severe deformation of peptidoglycan layer was observed after complete unfolding of BgaA.
